# BRD2 bridges TFIID and histone acetylation to promote transcriptional initiation

**DOI:** 10.1101/2025.09.20.676678

**Authors:** Bin Zheng, Ruxuan Qiu, Sarah Gold, Marta Iwanaszko, Yuki Aoi, Benjamin Charles Howard, Madhurima Das, Ali Shilatifard

**Author notes:** Correspondence to: Ali Shilatifard and/or Bin Zheng, Department of Biochemistry and Molecular Genetics, The Louis A. Simpson and Kimberly K. Querrey Biomedical Research Center, Room 7-515, 303 East Superior Street, Chicago, IL 60611, USA.

## Abstract

Members of the bromodomain and extraterminal domain (BET) protein family play a central role in transcription by RNA Polymerase II (Pol II). Small-molecule inhibitors that block interaction between BET bromodomains and acetylated histones have been developed to achieve therapeutic benefit. However, the BET protein BRD4 does not require bromodomains to perform its major transcriptional elongation function, and the mechanisms by which other BET proteins regulate transcription remain incompletely understood. Addressing the disparity between pan-BET degraders and BRD4-specific depletion, we report that the BET protein BRD2 generally functions to promote transcriptional initiation in a bromodomain-dependent manner at both promoters and enhancers in human cells. We demonstrate that BRD2 bromodomains preferentially bind to tetra-acetylated histones harboring MOF-mediated H4K16ac, while the BRD2 C terminal domain facilitates recruitment of TFIID. Our studies provide mechanistic insight into the distinct roles of BRD2 in transcriptional initiation through the recruitment of TFIID and BRD4 in transcriptional elongation through the recruitment of CDK9 and controlling proper regulation of gene expression.

## Introduction

The BET (bromodomain and extraterminal domain-containing) family proteins BRD2, BRD3, BRD4, and BRDT are essential, multifunctional transcription coactivators that have been proposed as therapeutic targets for a variety of diseases, including cancer^1–4^. The two tandem bromodomains present in all BET family proteins confer each with the capacity to bind histone acetylation, but these bromodomains differ in their binding affinities for specific histone residues, and BET proteins also contain other functional domains; together, these differences enable distinct functions for individual BET proteins, some of which have been previously reported^5–9^. Small molecule inhibitors developed to target BET bromodomains are highly potent disruptors of BET protein binding to histone acetylation, which is associated with transcriptionally active chromatin^10,11^. However, the disparities between the transcriptional and therapeutic outcomes of these BET inhibitors and those of targeted BET protein degraders such as proteolysis targeting chimeras (PROTACs)^12–14^ have highlighted the need to characterize distinct functions of individual BET proteins in order to support the development of effective, nontoxic therapeutic approaches that selectively target these distinct BET protein functions^15,16^.

In previous studies, we have used acute depletion strategies such as auxin-induced degradation (AID)^17,18^ to compare the genome-wide impacts of individual BET proteins on transcription in living cells^19,20^. We reported that BRD4 can perform its master transcriptional regulatory function, in which it facilitates the release of promoter-proximal paused RNA Polymerase II (Pol II) by the CDK9-containing complex PTEFb, independently of its bromodomains^20^. This finding indicates that contrary to their supposed mechanism of action, inhibitors designed to target BET bromodomains may fail to impede the major transcriptional regulatory function of BRD4.

Nevertheless, cytotoxicity and antitumor efficacy have been demonstrated for BET bromodomain inhibitors in preclinical studies, and there is evidence of clinical benefit in some patients^21–26^. This discrepancy led us to hypothesize that the apparent efficacy of BET bromodomain inhibitors could be at least partially due to their impact on BET proteins other than BRD4.

As the most abundant BET protein in nearly all cell types, BRD2 is an important factor involved in many essential processes. For example, BRD2 regulates chromatin structures by collaborating with CTCF or cohesin^27,28^, is essential for embryonic development^29^, and regulates erythroid gene expression in concert with BRD3^30^. BRD2 has also been proposed as a therapeutic target for cancer^31,32^. Our initial study of acute BRD2 depletion revealed that BRD2 is required for maintaining the presence of transcriptional machinery at BET-bound enhancers^19^, a finding that was confirmed by another group using a different method^33^. However, the mechanism by which BRD2 regulates transcription at enhancers and to what extent perturbing this function contributes to the efficacy of BET treatment remains largely unknown.

Studies in yeast have implicated the BET homologues Bdf1/2 in transcription initiation^34^. A study in mammalian cells also reported that BRD2 facilities transcription initiation at loci with low H3K4me3 levels via direct interaction with TAF3^35^, which would otherwise be recruited to loci with abundant H3K4me3^36,37^. However, recent studies indicate that H3K4me3 is not instructive for transcription initiation^38,39^. We therefore hypothesized that BRD2 might play a more general role to facilitate transcriptional initiation (rather than specifically compensating for lack of abundant H3K4me3), and we aimed to further investigate and elucidate the molecular mechanism by which BRD2 might do so.

In metazoans, the RNA Pol II transcription cycle requires participation from a series of transcription factors to coordinate the sequential steps of initiation, pausing, elongation, and termination^40,41^. Transcription initiation starts with the recognition of promoter regions by the general transcription factor TBP/TFIID, via both TATA box-dependent and -independent mechanisms, followed by formation of the preinitiation complex (PIC) with other essential general transcription factors including TFIIA, TFIIB, TFIIE, TFIIF, and TFIIH^42–45^. Initiation is completed upon promoter escape, when engaged Pol II begins to transcribe nascent RNA. After transcribing 20-120 nucleotides Pol II pauses at a promoter-proximal region, where elongation factors such as SPT5/6 and PAF1C are recruited before PTEFb (generally recruited by BRD4) releases Pol II into productive transcript elongation and then termination to finish the transcription cycle^46–48^. The less known BRD2’s potential involving in the initiation stage could be at the very beginning given the documented interaction of between BRD2 and several TFIID components^35,49^.

Here, using acute depletion and genetic complementation and biochemical and molecular methods, we demonstrate that BRD2 promotes Pol II transcription in a bromodomain-dependent manner at active transcription sites, including promoters and enhancers. We demonstrate that BRD2 specifically promotes transcription at sites with tetra-acetylated histone H4 harboring MOF-mediated H4K16ac, to which BRD2 bromodomains preferentially bind. Moreover, we demonstrate that BRD2 does so via interaction between its C terminal domain and TFIID. Our results support a model in which the major BET family proteins BRD2 and BRD4 fulfill non-redundant functions by regulating distinct steps within the Pol II transcription cycle.

## Results

### Disparate impacts of BET protein depletion suggest transcriptional regulatory roles for BRD2/BRD3 upstream of BRD4

Our previous studies addressing the disparity in Pol II occupancy profiles upon BET inhibition versus degradation led to the finding of bromodomain-independent BRD4 function in Pol II pause release and elongation control^19,20^. At the same time, we also observed by ChIP-seq that Pol II occupancy was lost from thousands of active transcription loci upon BET degradation, with the most profound impact at BET-bound enhancers^19^. To gain greater clarity regarding the impacts of BET protein depletion on Pol II, we carried out PRO-seq to analyze Pol II occupancy with high resolution upon pan-BET depletion (via dBET6 treatment) in WT DLD-1 cells, in comparison to BRD4-specific depletion (by auxin treatment) in BRD4-AID cells. Although Pol II occupancy within gene bodies was decreased by both treatments, consistent with the well-established function of BRD4 in Pol II pause release, there was a major difference in Pol II occupancy at promoters, which was increased by specific BRD4 depletion (consistent with Pol II pausing) but reduced by pan-BET depletion (Fig. 1a,b). Clustering Pol II-transcribed promoter regions by difference in Pol II distribution revealed that Pol II occupancy was reduced at nearly half of these promoters upon pan-BET depletion, while at the remaining promoters, increases in Pol II occupancy were strongly attenuated relative to specific BRD4 depletion (Fig. 1b). These observations were in line with published NET-seq data for similar conditions^50^, but the underlying mechanism was not clear. We speculated that depletion of another BET protein (BRD2 and/or BRD3) might “cancel” the BRD4 depletion-induced pausing effect (Extended Data Fig. 1a) if this BET protein functioned to regulate transcription prior to the promoter-proximal pausing step.

**Fig. 1:**
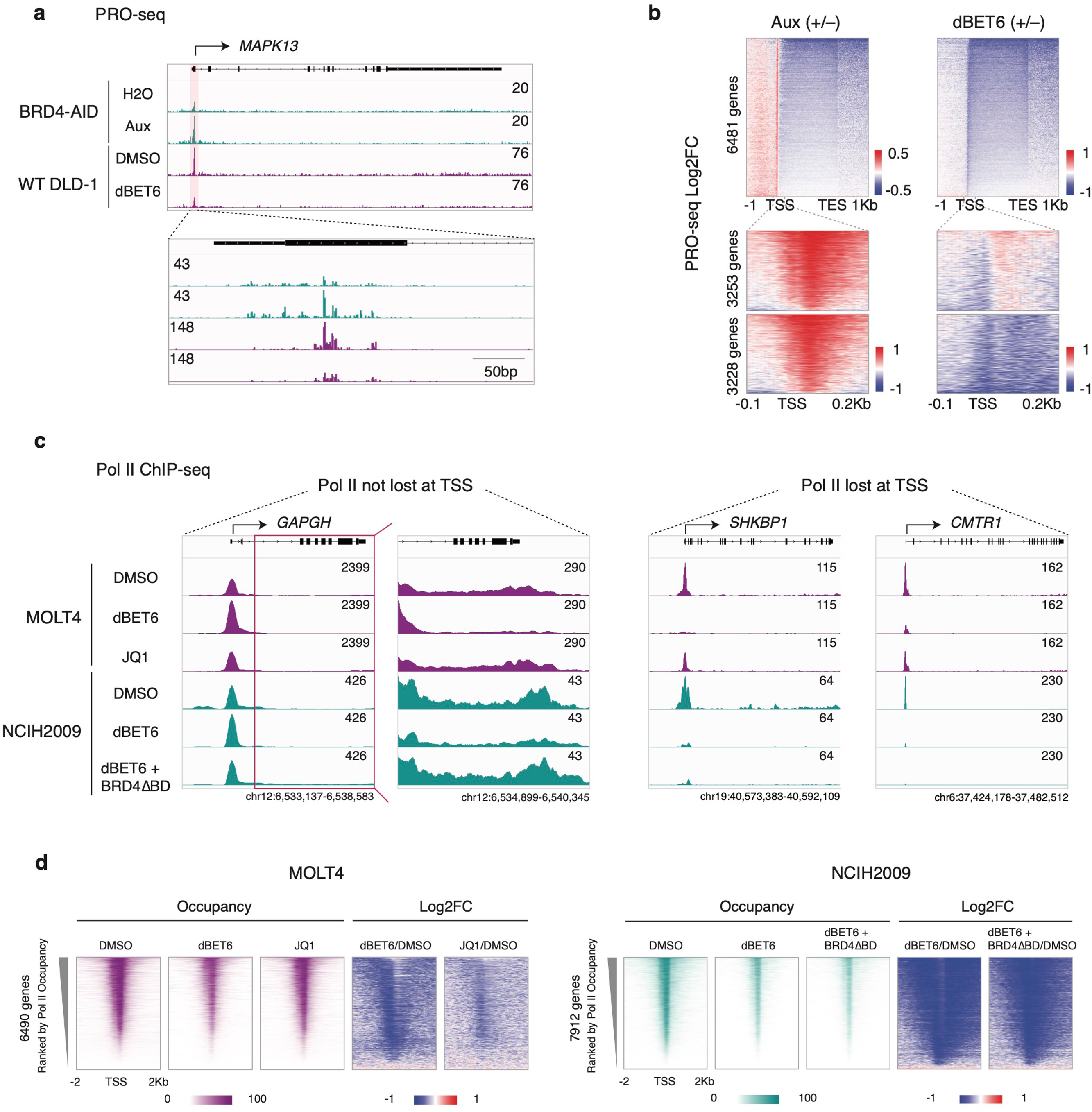
Distinct impacts of BET protein disruption strategies suggest transcriptional regulatory roles for BRD2/BRD3 upstream of BRD4. (A) Genomic track visualizations of PRO-seq signal at the *MAPK13* gene locus in BRD4-AID DLD-1 cells and in parental (WT) DLD-1 cells, with a zoom-in view of signal at the promoter region. BRD4-AID DLD-1 cells were treated with H2O or auxin (Aux, 500 uM) for 3h; WT DLD-1 cells were treated with DMSO or dBET6 (250 nM) for 2h. (B) Heatmap of Log2FC in PRO-seq signal at 6481 genes for the conditions described in panel a. A zoom in view of signal at promoter regions is also shown, with the datasets partitioned into two k-means clusters. (C) Genomic track visualizations of Pol II ChIP-seq signal in MOLT4 cells treated with DMSO, 100 nM dBET6 or 1 uM JQ1 for 2h, and in NCIH2009 cells treated with DMSO or dBET6 with or without overexpression of a BRD4 construct lacking bromodomains (BRD4ΔBD). Signals are visualized at the *GAPDH* gene loci, where Pol II occupancy was retained at the TSS upon dBET6 treatment, and at the *SHKBP1* and *CMTR1* gene loci, where it was lost. A zoom in view of Pol II signal within the gene body is also shown for *GAPDH.* Data for Pol II ChIP-seq in MOLT4 cells was downloaded from Winter et al., *Mol Cell*^12^; data for Pol II ChIP-seq in NCIH2009 cells was previously published^20^, but was realigned and normalized to spike-in control. (D) Heatmap of Pol II ChIP-seq signal and Log2FC fold change in signal for the conditions described in panel c, at 6490 and 7912 Pol II transcribed genes in MOLT4 and NICH2009 cells, respectively.

To confirm that the phenomenon of Pol II loss upon pan-BET depletion was not limited to DLD-1 cells, we re-analyzed the first published Pol II ChIP-seq of dBET6 treatment by Winter et al. in the MOLT4 T lymphoblast cell line^12^, as well as our previously published Pol II ChIP-seq of dBET6 treatment in the NCIH2009 cell line^20^, which was derived from lung adenocarcinoma. Data were also analyzed for MOLT4 cells treated with the BET bromodomain inhibitor JQ1 and for NCIH2009 cells that were induced to express a bromodomainless BRD4 construct (BRD4ΔBD) prior to treatment with dBET6. Consistent with what we observed in DLD-1 cells, there appeared to be two groups of genes upon dBET6 treatment: one group exhibited a high level of dBET6-induced pausing, with Pol II accumulated at the promoter and reduced in the gene body (e.g. *GAPDH*), while the other group exhibited severe loss of Pol II from promoters (e.g. *SHKBP1* and *CMTR1*) (Fig. 1c). Importantly, MOLT4 cells treated with JQ1 exhibited a similar pattern of Pol II loss from promoters (but to a lesser extent) as compared to cells treated with dBET6 (Fig. 1c,d). Additionally, though we have previously shown that expression of the bromodomainless BRD4 construct (BRD4ΔBD) could rescue Pol II pause release in NCIH2009 cells, BRD4ΔBD expression could only rescue genes with a high level of dBET6-induced pausing, but not at genes with severe Pol II loss (Fig. 1c,d and Extended Data Fig. 1b), indicating a mechanism upstream of the BRD4-regulated pause-release step.

### Disruption of bromodomain-dependent BRD2 chromatin binding permits formation of nuclear BRD2 protein assemblies

Since the expression level of BRD2 is much higher than that of BRD3 in DLD-1 cells (Extended Data Fig. 2a), we chose to focus on defining the transcriptional regulatory function of BRD2 and the impact of bromodomain inhibition on this function. We therefore generated Dox-inducible constructs to express GFP-tagged and nuclear-localized full length (FL) or bromodomain deletion mutant (ΔBD) BRD2 (Fig. 2a). We initially checked the nuclear localization of these GFP-tagged BRD2 constructs using fluorescence microscopy. Unexpectedly, GFP-tagged BRD2ΔBD (but not BRD2FL) formed distinct foci, or protein assemblies, in the nuclei (Fig. 2a). We then tested whether BET bromodomain inhibition could impart the same phenotype in cells expressing GFP-tagged BRD2FL. We observed similar BRD2FL foci at higher JQ1 doses, with apparent saturation at 10 uM JQ1 (Fig. 2b), suggesting that the displacement of BRD2 from chromatin is complete in cells treated with JQ1 at high doses. We also tested the impact of JQ1 treatment on the localization of endogenous BRD2 by immunofluorescent staining in other cell lines, which confirmed that this BET bromodomain deletion/inhibition-induced phenotype is neither an artifact of exogenous construct expression nor restricted to DLD-1 cells (Extended Data Fig. 2b-d).

**Fig. 2:**
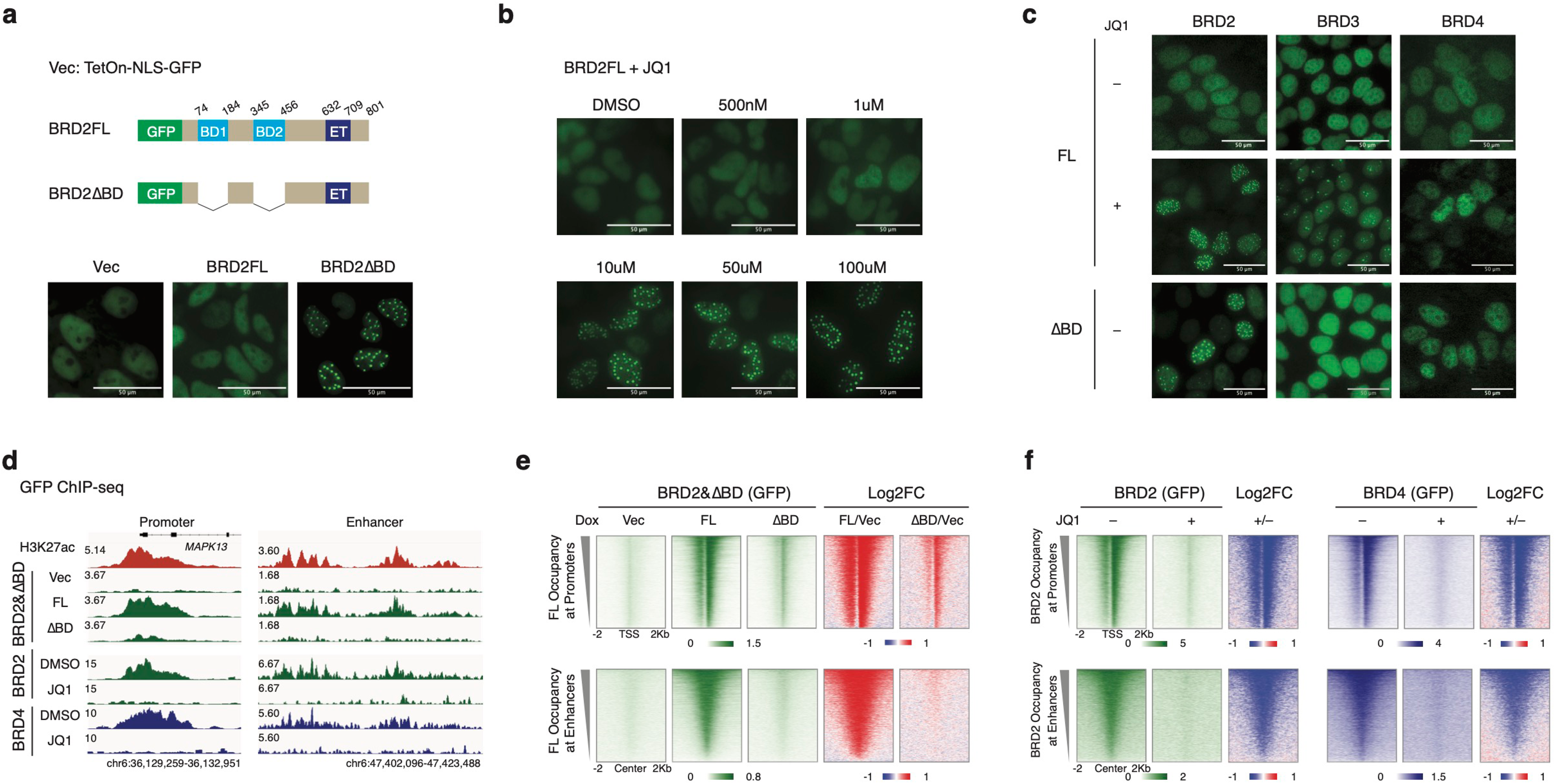
Disruption of bromodomain-dependent BRD2 chromatin binding permits formation of nuclear BRD2 protein assemblies. (A) Schematic illustration of full length (BRD2FL) and bromodomain deletion mutant (BRD2ΔBD) BRD2 constructs expressed from the TetOn-NLS-GFP vector (above) and representative fluorescence microscopy images of GFP signal in untreated BRD2-AID DLD-1 cells following Dox-induced expression (50 nM, 20h prior to imaging) of the vector control or BRD2 constructs (below). Nuclear foci are evident in cells expressing the GFP-BRD2ΔBD construct. Scale bar, 50 um. (B) Representative fluorescence microscopy images of GFP signal in the nuclei of DLD-1 cells expressing the GFP-BRD2FL construct and treated with DMSO or the BET bromodomain inhibitor JQ1 (100 nM, 1 uM, 10 uM, 50 uM, or 100 uM in DMSO) for 18h. Nuclear foci are evident upon JQ1 treatment at doses ≥10 uM. Scale bar, 50 um. (C) GFP signal in the nuclei of DLD-1 cells expressing GFP-tagged full length (FL) or bromodomain deletion (ΔBD) constructs for each BET protein family member. Representative images are shown for cells expressing GFP-BRD2, BRD3, or BRD4 FL constructs treated without (top row) or with 10 uM JQ1 over night (middle row), as well as cells expressing GFP-BRD2, BRD3, or BRD4 ΔBD constructs (bottom row). Nuclear foci are evident in JQ1-treated cells expressing the GFP-BRD2 or BRD3 FL constructs, and in untreated cells expressing the GFP-BRD2ΔBD construct. Scale bar, 50 um. (D) Genomic track visualizations of GFP ChIP-seq signal at representative promoter and enhancer (as indicated by H3K27ac ChIP-seq signal, top) in DLD-1 cells expressing the TetOn-NLS-GFP vector or the GFP-BRD2FL or GFP-BRD2ΔBD constructs, in DLD-1 cells expressing the GFP-BRD2FL construct and treated with DMSO or 10 uM JQ1 for 3h, and in BRD4-IAA7 DLD-1 cells expressing the GFP-BRD4FL construct and treated with DMSO or 10 uM JQ1 for 3h. H3K27ac ChIP-seq was performed in untreated BRD2-AID DLD-1 cells (true throughout the study). (E) Heatmaps of GFP ChIP-seq signal in DLD-1 cells expressing the TetOn-NLS-GFP vector or the GFP-BRD2FL or GFP-BRD2ΔBD constructs, with heatmaps of Log2FC in GFP signal relative to the vector control, at Pol II-transcribed gene promoters (n=6481) and at BET-bound enhancers (n=8147, as determined in previous work^19^; see Methods). (F) Heatmaps of GFP ChIP-seq signal in DMSO or JQ1-treated DLD-1 cells expressing the GFP-BRD2FL construct and in DMSO or JQ1-treated BRD4-IAA7 DLD-1 cells expressing the GFP-BRD4FL construct, with heatmaps of log2FC in GFP signal in JQ1-treated relative to DMSO-treated cells, at the promoters and enhancers described in panel e. Cells were treated as described in panel d.

To test whether the bromodomain disruption-induced formation of nuclear foci is common among BET family proteins, we generated additional constructs in the same TetOn-NLS-GFP vector to express GFP-tagged FL or ΔBD versions of BRD3 and BRD4. Cells expressing FL BRD2/3/4 were further treated with 10 uM JQ1. GFP-tagged BRD2 and BRD3 both formed nuclear foci upon JQ1 treatment, while BRD4 did not (Fig. 2c). However, only BRD2ΔBD formed these foci in the absence of JQ1 treatment (Fig. 2c). Moreover, JQ1-induced BRD3 foci were fewer and more spatially constrained than those of BRD2, consistent with a previous study demonstrating that JQ1 treatment re-localizes BRD3 to form dense foci in the nucleolus^49^.

Reasoning that the bromodomain disruption-induced formation of BRD2 foci must occur secondary to the displacement of BRD2 from chromatin, we performed GFP ChIP-seq to assess the impact of BET bromodomain disruption on BRD2 chromatin occupancy genome wide. GFP-BRD2FL occupancy was much higher than GFP-BRD2ΔBD occupancy at the promoters of Pol II-transcribed genes and at BET protein-bound enhancers (Fig. 2d). This apparent bromodomain dependence was particularly evident at enhancers, whereas promoters retained some BRD2ΔBD peaks (Fig. 2d,e), which could potentially indicate bromodomain-independent chromatin association for BRD2 at promoters^51^. These retained BRD2ΔBD peaks appeared more obvious than those observed for BRD4ΔBD in our previous study (Fig. 2e and Extended Data Fig. 2e)^20^. There was no evidence of BRD2ΔBD relocalization. GFP ChIP-seq also indicated that for both GFP-BRD2 and GFP-BRD4, occupancy was efficiently abolished by JQ1 treatment at 10uM (Fig. 2d,f). Together with fluorescence microscopy imaging, the ChIP-seq analysis of GFP-tagged BRD2 occupancy indicates that BET bromodomain inhibition induces a transition from histone acetylation-bound BRD2 to chromatin-unbound nuclear BRD2 protein assemblies.

Higher-order protein assemblies can take a variety of forms, including aggregates and condensates. In murine embryonic stem cells, researchers were previously able to image BRD4-containing co-activator condensates at super-enhancers, as well as JQ1-disruptable condensates in which Pol II and Mediator interact^52,53^. We observed the formation (rather than disruption) of BRD2 foci upon treatment with JQ1, indicating that these BRD2 foci are different from previously observed BRD4 condensates. Moreover, the fact that the foci form concomitant to displacement of BRD2 from chromatin is similar to/consistent with phenomena such as the assembly of PolyQ containing proteins into aggregate foci upon disruption of their DNA binding, when they are displaced from chromatin and lose the benefit of its solubilizing effect^54^.

### BRD2 promotes Pol II transcription initiation at promoters and enhancers

To understand the transcriptional impact of the BET bromodomain inhibition-induced transition from histone acetylation-bound BRD2 to chromatin-unbound BRD2 protein assemblies, we first carried out ChIP-seq for Pol II chromatin occupancy upon BRD2 depletion and rescue via expression of either BRD2FL or BRD2ΔBD (Fig. 3a,b). Acute depletion of endogenous BRD2 resulted in a genome-wide reduction of Pol II occupancy at both promoters and enhancers, which was fully restored by the expression of BRD2FL but not BRD2ΔBD (Fig. 3b,c and Extended Data Fig. 3a). BRD2ΔBD did partially restore Pol II occupancy, presumably due to its residual, potentially bromodomain-independent binding (Fig. 2e); however, this rescue was severely impaired relative to BRD2FL. As expected, JQ1 treatment with and without subsequent washout phenocopied endogenous BRD2 depletion with and without prior expression of BRD2FL (Fig. 3b,c and Extended Data Fig. 3a). In contrast to acute BRD2 depletion, we previously demonstrated that acute depletion of endogenous BRD4 results in genome-wide Pol II pausing that can be rescued by the expression of either BRD4FL or BRD4ΔBD (Extended Data Fig. 3b). These data indicate that chromatin-associated BRD2 promotes Pol II occupancy at active transcription sites across the genome, that it does so primarily via its bromodomains, and that it may also do so via unknown bromodomain-independent mechanisms. Additionally, these data suggest that the JQ1-induced loss of Pol II from promoters and enhancers is attributable to BRD2.

**Fig. 3:**
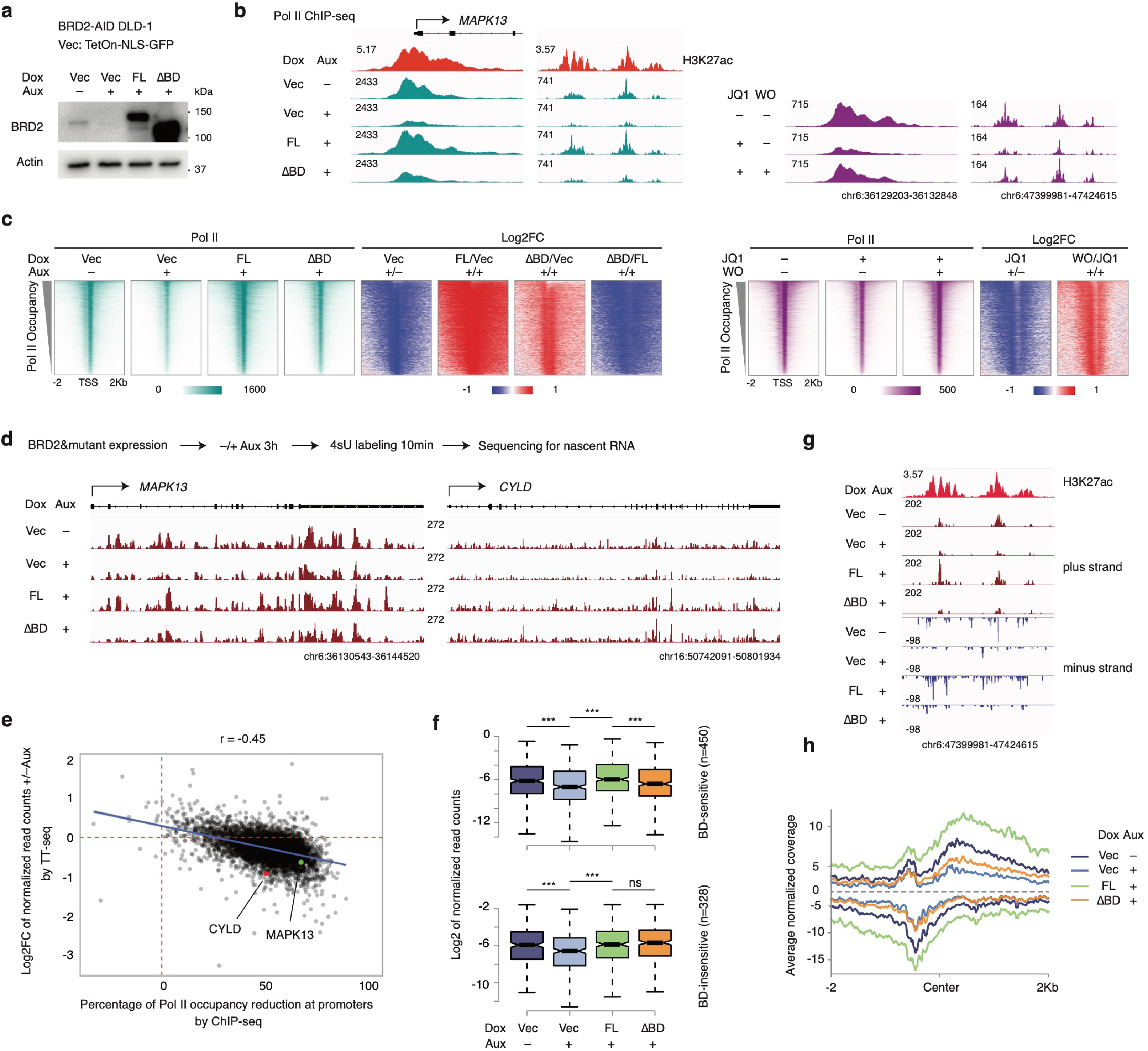
BRD2 promotes Pol II transcription initiation at promoters and enhancers. (A) Western blot analysis of endogenous BRD2 and/or construct protein expression levels in untreated BRD2-AID DLD-1 cells, upon endogenous BRD2 depletion in auxin-treated BRD2-AID DLD-1 cells, and in auxin-treated BRD2-AID DLD-1 cells expressing the GFP-BRD2 FL or ΔBD constructs (as shown in Fig. 2a). BRD2-AID DLD-1 cells were treated with H2O or auxin (500 uM) for 3h, and BRD2 construct expression was induced via 50 nM Dox treatment for 2 days. Blots were probed for BRD2, with actin as a loading control. (B) Genomic track visualizations of Pol II ChIP-seq signal at representative promoter and enhancer in untreated and auxin-treated BRD2-AID DLD-1 cells expressing the TetOn-NLS-GFP vector, auxin-treated BRD2-AID DLD-1 cells expressing the GFP-BRD2FL or GFP-BRD2ΔBD constructs, and DLD-1 cells expressing the GFP-BRD2FL construct that were additionally treated with DMSO for 3h (-/-) or with 10 uM JQ1, for 3h (+/-) or for 3h followed by washout for 6h (+/+). (C) Heatmaps of Pol II ChIP-seq signal at the promoters of genes actively transcribed by Pol II (n=6481), with log2FC in Pol II signal in the indicated comparisons, for the conditions described in panel b. (D) Workflow for TT-seq analysis of BRD2 depletion and rescue via expression of GFP-BRD2FL or GFP-BRD2ΔBD, shown with genomic track visualization of TT-seq signal at the example genes *MAPK13* and *CYLD*, which are downregulated upon BRD2 depletion. TT-seq analysis was performed on untreated and auxin-treated BRD2-AID DLD-1 cells expressing the TetOn-NLS-GFP vector, and on auxin-treated BRD2-AID DLD-1 cells expressing the GFP-BRD2FL or GFP-BRD2ΔBD constructs. (E) Scatter plot showing the relationship between changes in nascent transcription and Pol II promoter occupancy upon BRD2 depletion. The Y-axis represents the log2 FC in TT-seq signal (normalized read counts) within gene bodies (from TSS to TES) for auxin-treated vs untreated BRD2-AID DLD-1 cells. The X-axis represents the percentage of reduction in Pol II ChIP-seq signal density at promoters (±1kb flanking TSS). *MAPK13* and *CYLD* (featured in panel d) are annotated. (F) Boxplot showing the log2 transformed normalized TT-seq read counts in the gene body of BRD2-dependent genes, for each of the BRD2 depletion and rescue conditions described in panel d. BRD2-dependent genes (N=1883) are divided into BD-sensitive (N=450) and BD-insensitive (N=328) groups. BD-sensitive genes were selected as log2(ΔBDvsFL) <= −0.415, while BD-insensitive genes were selected as log2(ΔBDvsFL) >=0. (***: p-value < 0.001, ns: p-value > 0.2, Wilcoxon test). (G) Genomic track visualization of TT-seq signal at a representative enhancer (as indicated by H3K27ac ChIP-seq signal) for the rescue experiment described in panel d. TT-seq signal is shown for both plus and minus strands. (H) Strand-specific metagene plot of TT-seq signal at BET-bound enhancers (n=8147) for the rescue experiment described in panel d.

Decreased Pol II occupancy at promoters might reflect impaired transcription initiation, or it could reflect potentiated release of promoter-proximal paused Pol II. To distinguish between these possibilities, we characterized the transcriptional impact of depleting histone acetylation-bound BRD2 directly by performing TT-seq to assess nascent transcription in endogenous BRD2-depleted cells expressing either BRD2FL or BRD2ΔBD (Fig. 3d). We observed a trend in which the more severe the loss of Pol II ChIP-seq signal from promoters upon BRD2 depletion, the more that TT-seq signals were reduced (Fig. 3e). Under a less stringent Log2FC cutoff (0.415), we observed many more downregulated (N=1883) than upregulated (111) genes upon BRD2 depletion (Extended Data Fig. 3c), consistent with the interpretation that genome-wide loss of Pol II from promoters mainly reflects a transcription initiation defect in BRD2-depleted cells. Among the genes downregulated upon BRD2 depletion, we identified a group of genes for which expression was fully rescued by BRD2FL but not BRD2ΔBD (BD-sensitive, N=450) and another group of genes (BD-insensitive, N=328) that could be rescued by either BRD2FL or BRD2ΔBD (Fig. 3d,e and Extended Data Fig. 3d). Re-analysis of ChIP-seq data for Pol II occupancy at genes in these two groups was consistent with the TT-seq analysis (Extended Data Fig. 3e). Although no significantly enriched pathways were found for the BD-insensitive genes, BD-sensitive genes were enriched for several important pathways such as Wnt signaling and embryonic morphogenesis, reflecting the known role of BRD2 in embryonic development^55,56^ (Extended Data Fig. 3f). TT-seq also revealed loss of nascent transcription at enhancers upon BRD2 depletion, which is fully rescued by BRD2FL but not BRD2ΔBD (Fig. 3g,h), indicating that BRD2 also promotes transcription initiation at enhancers. Consistent with Pol II ChIP-seq data, the observed impact of BRD2 depletion on nascent transcription was more dramatic at enhancers than promoters.

Taken together, these Pol II ChIP-seq and TT-seq analyses indicate that BRD2 plays a primary role in promoting transcription by RNA Pol II at promoters and enhancers that differs from the role of BRD4 in transcriptional elongation control. They also indicate that unlike BRD4, BRD2’s bromodomains are generally required for its transcription-promoting function. Moreover, they suggest that BRD2 mediates transcription initiation.

### The BRD2 C terminus facilitates TFIID recruitment

Given the potential function of BRD2 in transcription initiation, we were curious about the BRD2 interactome. We initially carried out GFP IP-MS for GFP-BRD2FL with/without JQ1 or GFP-BRD2ΔBD. For cells expressing GFP-BRD2FL, we detected interaction with TBP and associated factors belonging to the TFIID initiation complex in all six replicates (Fig. 4a), consistent with BRD2 interactome data from previous studies^35,49^. However, in contrast to a previous report that BRD2 interacts with TBP via its first bromodomain^57^, we also detected these BRD2-TFIID interactions when bromodomains were deleted or inhibited via JQ1 treatment (Fig. 4a and Extended Data Fig. 4a). In the DepMap (The Cancer Dependency Map Project at Broad Institute: https://depmap.org/portal) database, genes encoding components of the TFIID complex are among those with the greatest positive correlation with BRD2 for CRISPR impact across all available cell lines (Fig. 4b and Extended Data Fig. 4b). We also interrogated interactions between endogenous BRD2 and TFIID subunits by generating a new BRD2-IAA7 degron line (E2) (Fig. 4c) that would allow for direct comparison of endogenous, V5-tagged BRD2 with the endogenous, V5-tagged BRD4 in a BRD4-IAA7 degron line (G3) that we previously generated in the same DLD-1 background^20^. IP for the V5 tag confirmed that endogenous BRD2 interacts with the TFIID subunits TBP and TAF1, while BRD4 interacts with the well-characterized PTEFb complex (Fig. 4d), confirming distinct transcriptional regulatory roles for BRD2 and BRD4.

**Fig. 4:**
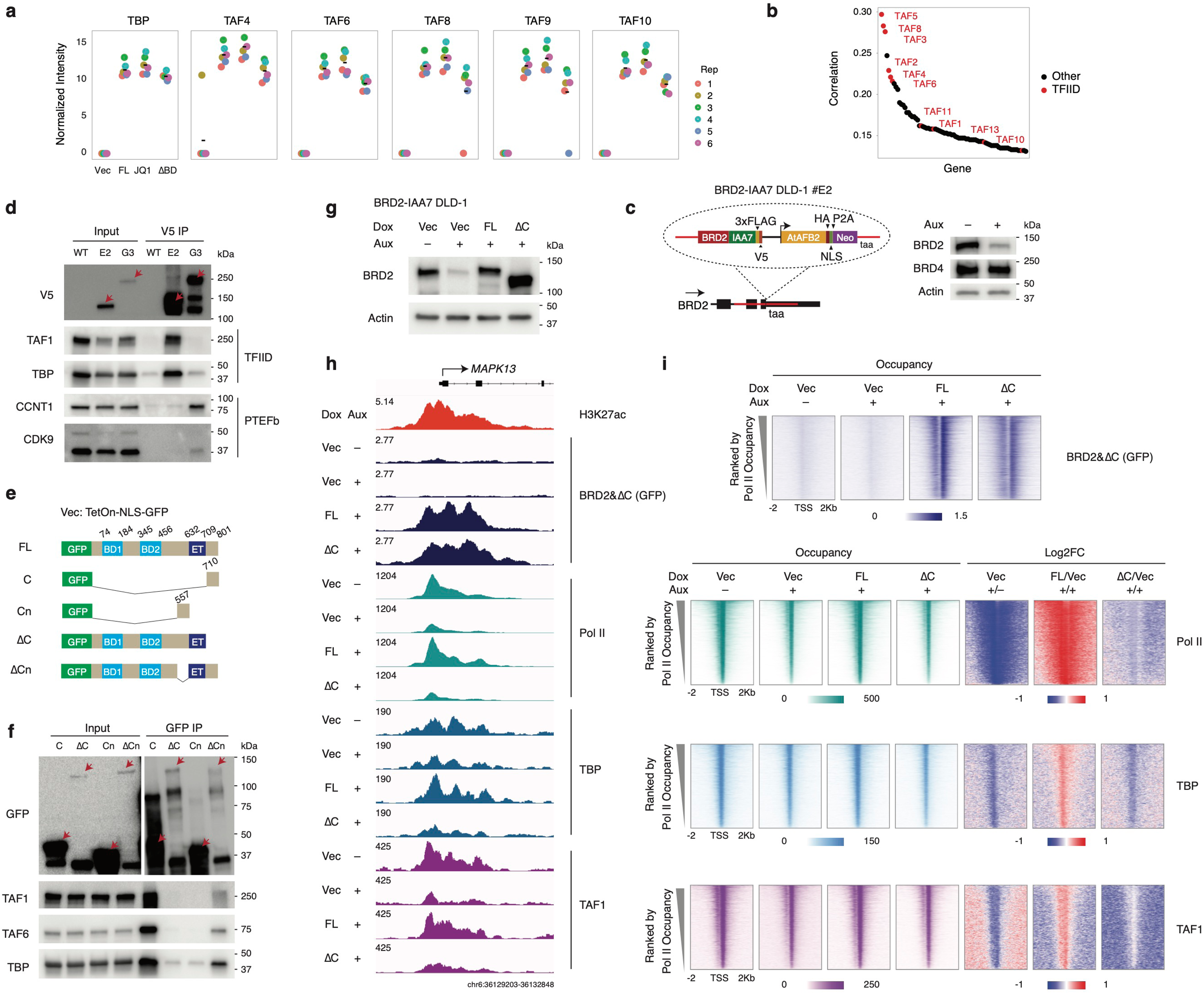
The BRD2 C terminus facilitates TFIID recruitment. (A) GFP IP-MS from WT DLD-1 cells expressing the TetOn-NLS-GFP vector, the GFP-BRD2FL construct (treated with DMSO or 10 uM JQ1), or the GFP-BRD2ΔBD construct. (B) Ranked plot of top 100 genes for which CRISPR effect score is most positively correlated with that of *BRD2* across all cell lines available from DepMap. Genes encoding TFIID subunits are named and highlighted in red. (C) Schematic illustrating the design of the BRD2-IAA7 degron line, similar to that of the BRD4-IAA7 degron line reported in our previous study^20^, with western blot validating the efficient and specific degradation of BRD2 (but not BRD4) upon auxin treatment for 3h in clone #E2. (D) Western blot of V5 IP for endogenous BRD2 or BRD4 in BRD2-IAA7 (E2) or BRD4-IAA7 (G3) DLD-1 cells, respectively. Blots were probed for the V5 tag, the TFIID subunits TBP and TAF1, and the PTEFb subunits CCNT1 and CDK9. Red arrows indicate the target bands. (E) Schematic illustration of the BRD2FL construct (also shown in Fig. 2a) and BRD2 mutant constructs expressed from the TetOn-NLS-GFP vector. C: Fragment at the C terminus of BRD2 containing the coiled-coil domain, immediately C terminal to the ET domain. Cn: BRD2 fragment immediately N terminal to the ET domain. ΔC: BRD2 with C deletion. ΔCn: BRD2 with Cn deletion. (F) Western blot of GFP IP for the BRD2 mutant constructs illustrated in panel e. Blots were probed for GFP and for the TFIID subunits TAF1, TAF6, and TBP. For better visualization of target bands, a long and short exposure are shown for input and for IP, respectively. Red arrows indicate the bands corresponding to the GFP-tagged mutants. (G) Western blot analysis of endogenous BRD2 depletion by auxin treatment for 3h in BRD2-IAA7 DLD-1 cells and the expression levels of GFP-tagged BRD2 FL and ΔC constructs (illustrated in panel e) induced by Dox treatment in BRD2-depleted cells. Blots were probed for BRD2 with actin as loading control. (H) Genomic track visualization of GFP, Pol II, TBP, and TAF1 ChIP-seq signals at the *MAPK13* locus in auxin-untreated/treated BRD2-IAA7 DLD-1 cells expressing the TetOn-NLS-GFP vector and in auxin-treated BRD2-IAA7 DLD-1 cells expressing the GFP-tagged BRD2 FL or ΔC constructs. GFP ChIP-seq was carried out after Dox treatment for 2 days, while ChIP-seq for Pol II, TBP, and TAF1 was carried out after Dox treatment for 36h in the same cells. (I) Heatmaps of GFP, Pol II, TBP, and TAF1 ChIP-seq signals at Pol II-transcribed gene promoters (n=6481) for the conditions described in panel h, with heatmaps of log2FC in Pol II, TBP, and TAF1signal upon of auxin depletion and upon rescue with the BRD2 FL or ΔC constructs.

Next, we sought to identify the BRD2 region responsible for TFIID binding. We generated a series of GFP-tagged BRD2 domain deletion mutant constructs (Extended Data Fig. 4c) and carried out GFP IP from construct-expressing cells, which narrowed down the binding site to the regions flanking the ET domain (Extended Data Fig. 4d). We then generated GFP-tagged constructs in which these regions were isolated (C and Cn) or deleted (ΔC and ΔCn) (Fig. 4e). GFP IP from cells expressing this second set of constructs revealed that the TFIID binding site is located in the C terminus of BRD2, which contains a predicted coiled-coil domain that is conserved from yeast to human and is essential for cell growth as demonstrated in murine erythroid progenitor cells^58^; deletion of this region (ΔC) abolished BRD2 binding to TFIID (Fig. 4f). Finally, after verifying the efficient depletion of endogenous BRD2 from chromatin in auxin-treated BRD2-IAA7 cells and impact on Pol II comparable to that seen in auxin-treated BRD2-AID cells (Extended Data Fig. 4e-g), we carried out the BRD2 depletion and rescue experiment in BRD2-IAA7 cells with the ΔC mutant (Fig. 4g). While the deletion of the C terminus domain in the ΔC mutant does not eliminate its chromatin binding, the ability of the ΔC mutant to rescue TBP and TAF1 recruitment, as well as Pol II occupancy, is dramatically decreased compared to that of full-length BRD2 (Fig. 4h,i and Extended Data Fig. 4h,i).

### Loss of Pol II and TFIID from chromatin upon BET inhibition is correlated with loss of BRD2

Given that 1) BRD2 depletion led to a loss of both Pol II and TFIID from chromatin, that 2) treatment with the BET inhibitor JQ1 phenocopies the impact of BRD2 depletion on Pol II occupancy, and that 3) JQ1 treatment has little impact on the bromodomain-independent function of BRD4 in Pol II pause release, we speculated that the major mechanism underlying transcriptional suppression upon JQ1 treatment might be the displacement of TFIID from chromatin, together with BRD2. To address this possibility, we tested the impact of BET inhibition on the BRD2/TFIID axis by treating WT DLD-1 cells with 10uM JQ1, followed by ChIP-seq analysis of chromatin occupancy for BRD2, Pol II, TBP, and TAF1 (Fig. 5a and Extended Data Fig. 5a). We observed an apparent pattern in which the more severe the loss of BRD2 from an active transcription site, the more dramatic the decrease in TBP and TAF1 binding at those sites, reflected in a greater impact on Pol II (Fig. 5a and Extended Data Fig. 5b,c). Our previous promoter and enhancer criteria excluded certain transcription loci. To be more inclusive, we called peaks in the control condition and overlapped BRD2, Pol II, TBP, and TAF1 binding sites, which resulted in identification of 23514 shared sites reflecting active Pol II transcription potentially regulated by the BRD2/TFIID axis (Fig. 5b). Plotting BRD2, Pol II, TBP, and TAF1 occupancy before and after JQ1 treatment together for these shared binding sites revealed an apparent correlation between loss of TFIID/Pol II and loss of BRD2 from chromatin upon bromodomain inhibition (Fig. 5c). To statistically evaluate the correlation, we computed the log2FC of Pol II and TBP/TAF1 occupancy upon JQ1 treatment at each of the binding sites and plotted against the log2FC of BRD2 occupancy. We found strong correlations between BRD2 loss and loss of TBP, TAF1 and Pol II from the 23514 shared binding sites but not from the 67305 control sites bound only by Pol II (Fig. 5d and Extended Data Fig. 5d). Consistent with both bromodomain-dependent and -independent BRD2 function (Fig. 3), JQ1 treatment has greater impact on Pol II occupancy at BD-sensitive than at BD-insensitive genes (Extended Data Fig. 5e).

**Fig. 5:**
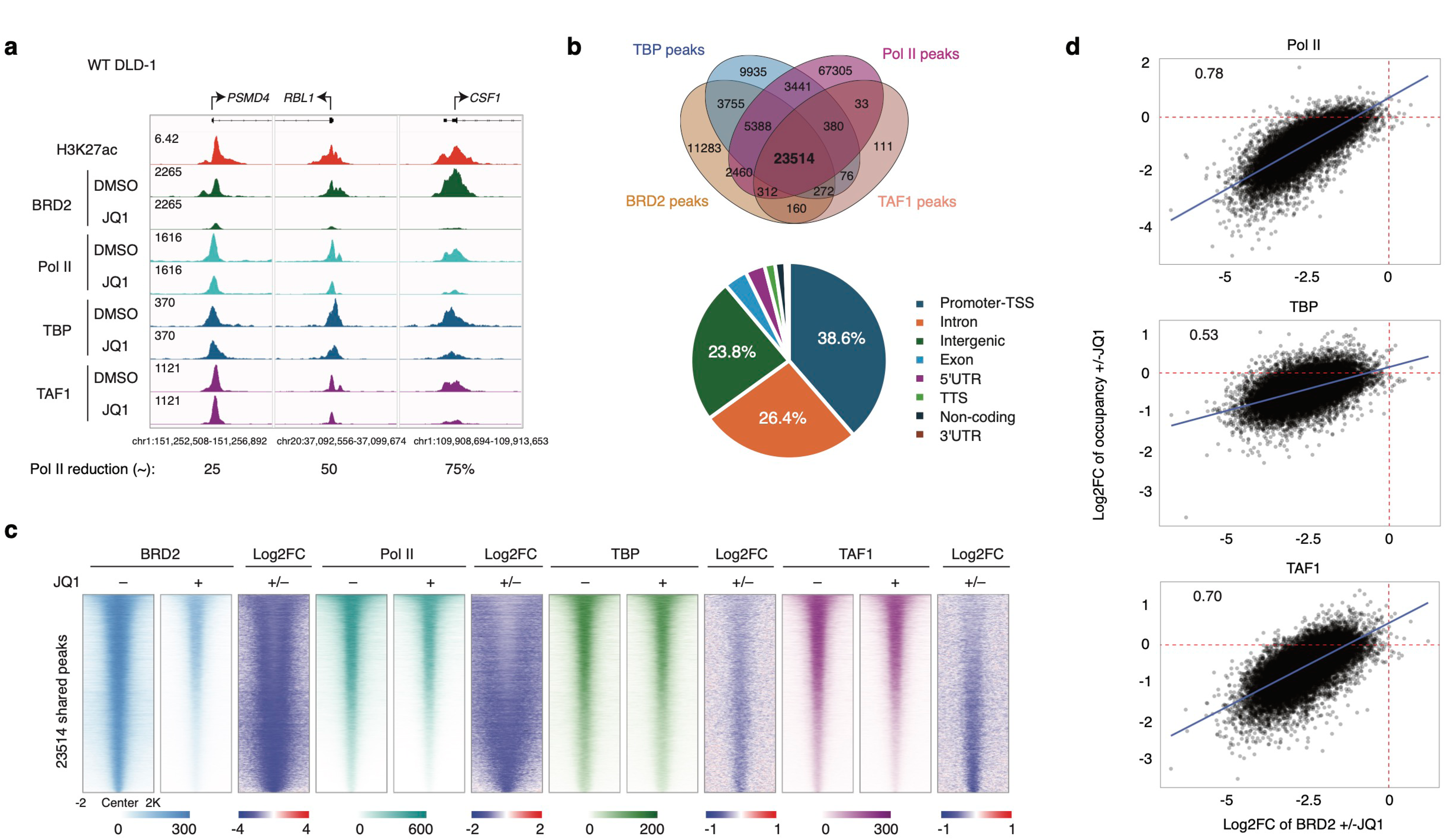
Loss of Pol II and TFIID from chromatin upon BET inhibition is correlated with loss of BRD2. (A) Genomic track visualization of BRD2, Pol II, TBP, and TAF1 ChIP-seq signals at the representative *PSMD4*, *RBL1*, and *CSF1* promoters in DLD-1 cells treated with DMSO or 10 uM JQ1 for 3h. (B) Overlap of BRD2, Pol II, TBP, and TAF1 ChIP-seq peaks in the DMSO-treated DLD-1 cells and annotation of the 23514 shared peaks bound by each factor. (C) Heatmaps of BRD2, Pol II, TBP, and TAF1 ChIP-seq signal at the 23514 shared peak regions identified in panel b, in DLD-1 cells treated with DMSO or 10 uM JQ1 for 3h, with heatmaps of log2FC in signal upon JQ1 treatment shown for each factor. (D) Scatter plot showing the correlation between the log2FC in BRD2 ChIP-seq signal density upon JQ1 treatment with those of Pol II, TBP, and TAF1 at the 23514 shared peak regions identified in panel b. Pearson correlation scores are shown.

### BRD2 and BRD4 BET bromodomains are nonfungible

BRD3 also interacts with TFIID (Extended Data Fig. 6a), and has long been implicated as functionally redundant with the structurally similar BRD2^6^. Because of the structural and functional similarities among BET family proteins, we sought to determine whether and to what extent BRD3 or BRD4 could functionally compensate for BRD2 loss. To do so, we repeated the BRD2 depletion and complementation/rescue experiment using full-length BRD2, BRD3, and BRD4 constructs as well as additional BRD2xBRD4 bromodomain exchange mutants (BRD4x2BD and BRD2x4BD), all generated in the TetOn-NLS-GFP vector (Fig. 6a,b and Extended Data Fig. 6b). Pol II ChIP-seq confirmed that BRD3 (but not BRD4) was able to functionally compensate for loss of endogenous BRD2, and also demonstrated that neither BRD2x4BD nor BRD4x2BD was able to rescue Pol II occupancy, consistent with a functional requirement for BRD2 domains other than its bromodomains (Fig. 6c,d and Extended Data Fig. 6c). Interestingly, BRD4x2BD and BRD2x4BD chromatin binding was largely impaired compared to that of WT BRD2 (Extended Data Fig. 6d,e), which may partially explain the inability of either mutant to rescue the effects of endogenous BRD2 depletion on Pol II. Additionally, ChIP-seq for GFP and BRD4 revealed mild decreases in the chromatin occupancy of both overexpressed and endogenous BRD4 upon BRD2 depletion (Extended Data Fig 6f,g), which may reflect a decrease of the Pol II-interacting layer of BRD4 (following loss of Pol II occupancy)^20,59^. Together, these data support redundant roles for BRD2 and BRD3 in transcriptional regulation, distinct from that of BRD4.

**Fig. 6:**
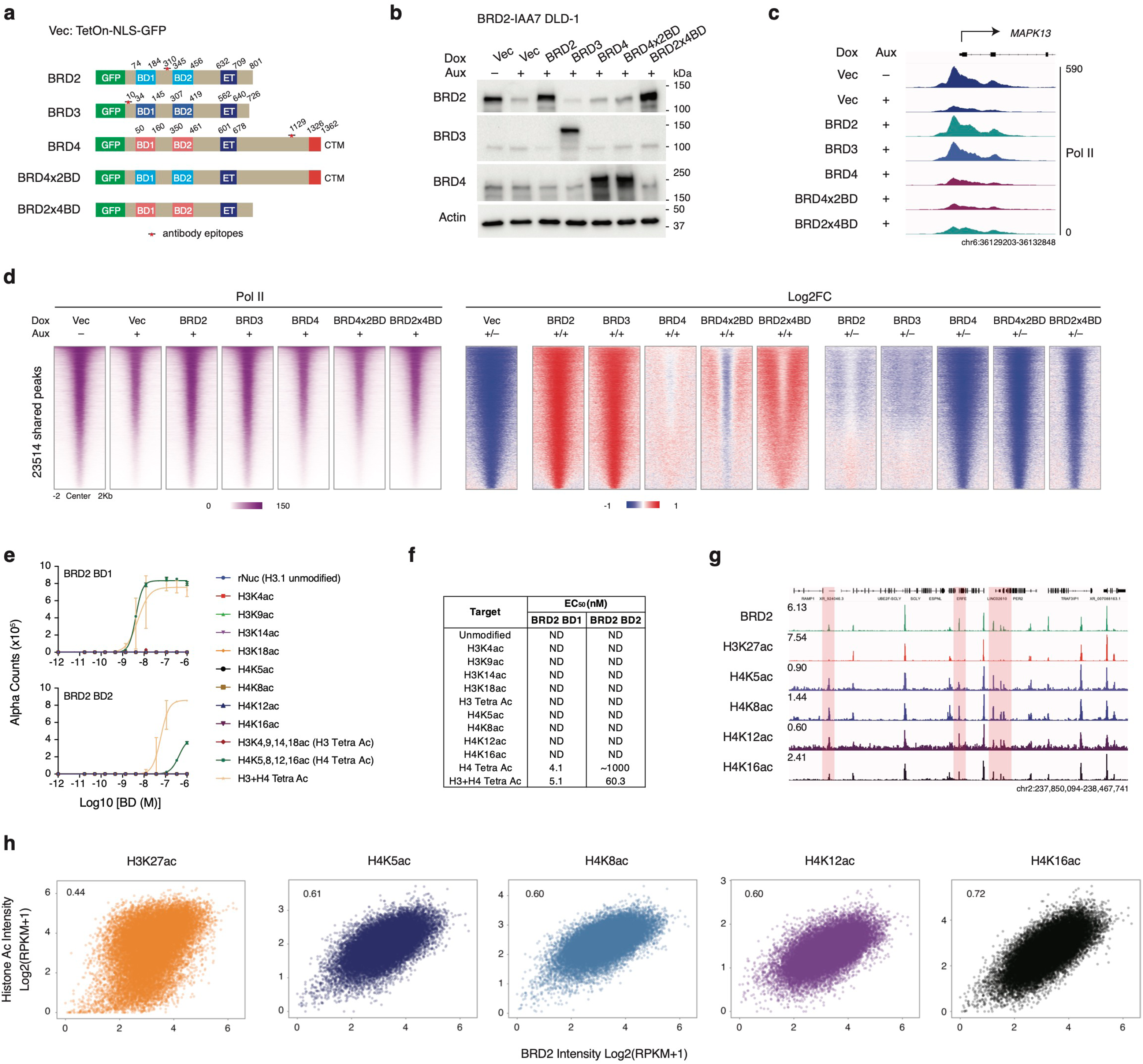
BRD2 and BRD4 bromodomains are nonfungible and BRD2 preferentially binds to tetra-acetylated histone H4 tails harboring H4K16ac. (A) Schematic illustration of BRD2, BRD3, BRD4 constructs (all full length) and domain swap constructs expressed from the TetOn-NLS-GFP vector. BRD4x2BD: BRD4 with bromodomains swapped for those of BRD2, BRD2x4BD: BRD2 with bromodomains swapped with those of BRD4. (B) Western blot of endogenous BRD2 depletion by auxin treatment and the expression of the constructs illustrated in panel a. Because 3xFLAG and V5 tags follow the IAA7 tag in BRD2-IAA7 cells (Fig. 4c), endogenous BRD2 is similar in size to the overexpressed GFP-BRD2FL construct. (C) Genomic track visualization of Pol II ChIP-seq signal at the *MAPK13* promoter in untreated BRD2-IAA7 DLD-1 cells and in auxin-treated BRD2-IAA7 DLD-1 cells expressing the TetOn-NLS-GFP vector or the GFP-tagged BET protein or domain swap constructs illustrated in panel a. (D) Heatmaps Pol II ChIP-seq signal and Log2FC in signal at the 23514 shared peak regions (bound by BRD2, Pol II, TBP, and TAF1) identified in Fig. 5b, for the BRD2 depletion and rescue conditions described in panel c. Log2FC heatmaps are shown for comparisons of construct-expressing cells to vector-expressing cells both with and without endogenous BRD2 depletion. (E) Assays for the binding affinity of GST-tagged BRD2 BD1 and BD2 domains to recombinant nucleosomes with different histone acetylation states. Nucleosome binding affinity assays were carried out by Epicypher. (F) EC_50_ for BRD2 BD1 and BD2 binding to recombinant nucleosomes with different histone acetylation states, as determined via the assays shown in panel e. (G) Genomic track visualization of ChIP-seq signals for BRD2, H3K27ac, H4K5ac, H4K8ac, H4K12ac, and H4K16ac over a representative region in BRD2-IAA7 DLD-1 cells. Loci where the pattern of BRD2 occupancy is similar to that of H4 acetylation (but not H3K27ac) are highlighted. (H) Scatter plots showing the correlations between BRD2 ChIP-seq signal intensity and those of H3K27ac, H4K5ac, H4K8ac, H4K12ac, and H4K16ac at the 23514 shared peak regions (identified in Fig. 5b), in BRD2-IAA7 DLD-1 cells. Pearson correlation scores are shown in the upper left-hand corner of each plot. ChIP-seq signal intensity was calculated as Log2 (RPKM+1) and normalized to total mapped reads and peak region length.

### BRD2 preferentially binds to tetra-acetylated histone H4 tails harboring H4K16ac

Given the finding that BRD2 and BRD4 bromodomains are nonfungible, we investigated the preferred histone acetylation binding targets for BRD2 bromodomains via recombinant bromodomain expression and in vitro nucleosome binding assays (Extended Data Fig. 6h). Neither bromodomain showed binding affinity for the mono acetylation that we tested, but strong binding was detected between either of the bromodomains and tetra-acetylated histone H4 tails (acetylated at lysines 5, 8, 12, and 16) (Fig. 6e,f). The binding appears to be specific, as neither bromodomain showed binding affinity for tetra-acetylated histone H3 without the additional presence of tetra-acetylated histone H4 (Fig. 6e,f). Interestingly, the first bromodomain of BRD2 (BD1) exhibits > 200-fold higher binding affinity to tetra-acetylated H4 than the second bromodomain (BD2) (Fig. 6e,f). To begin to test these findings in cells, we first compared BRD2 occupancy to that of H4K5, 8, 12, and 16ac or H3K27ac via ChIP-seq in DLD-1 cells. We observed BRD2 colocalization with these H4 acetylation marks even at loci lacking H3K27ac (Fig. 6g). Across the 23514 shared binding sites bound by BRD2, TFIID, and Pol II, we observed a stronger correlation between BRD2 occupancy and all four H4 acetylation marks than with H3K27ac (r=0.44), with highest correlation between BRD2 occupancy and H4K16ac (r=0.72) (Fig. 6h). H4K16ac is primarily deposited by the histone acetyltransferase MOF (also known as MYST1 or KAT8; throughout this manuscript, the protein is referred to as MOF and the encoding gene as *KAT8*)^60–62^. Taken together, these data suggest that the presence or absence of tetra-acetylated histone H4 harboring H4K16ac could be a determinant of BRD2 chromatin occupancy, and by extension, the chromatin occupancy of Pol II.

### Loss of MOF-mediated H4K16ac leads to decreased BRD2 and Pol II occupancy

To investigate whether there is a causal link between the presence of H4K16ac and BRD2 chromatin binding, we first carried out MOF knock down via CRISPR using sgRNA to target the MOF-encoding gene *KAT8*, which resulted in a reduced level of H4K16ac (Extended Data Fig. 7a). We observed a genome-wide reduction of BRD2 and Pol II occupancy at promoters in sgKAT8 relative to sgCtrl cells (Extended Data Fig. 7b,c). We then generated a MOF-dTAG^63^ cell line in the same BRD2-IAA7 background to avoid potential secondary effects of CRISPR knockdown (Fig. 7a). MOF was readily depleted within 1-2 hours following dTAG13 treatment (Fig. 7b). Bulk H4K16ac levels decreased more gradually, with notable decrease at 6-9 hours (two H4K16ac antibodies were tested), and some reduction in bulk H4K5ac and H4K8ac levels was also apparent (Fig. 7b), consistent with the known functions of the distinct MOF complexes MSL and NSL^61^. We then carried out ChIP-seq for H4K16ac, BRD2, and Pol II to assess the impact of MOF depletion in MOF-dTAG cells treated with dTAG13 for 9h, using antibodies from two different manufacturers (CST and Abcam) to detect H4K16ac. ChIP-seq analysis revealed a significant reduction in H4K16ac upon MOF depletion, especially across genebodies, whereas H4K16ac was retained at many promoters (Fig. 7c,d). We observed minimal changes in BRD2 and Pol II occupancy at loci that retained H4K16ac upon MOF depletion, in contrast to a severe loss of both BRD2 and Pol II at loci where H4K16ac was diminished (Fig. d,e). We calculated the Log2FC of BRD2 and Pol II intensity at promoter regions for the 8461 Pol II transcribed genes upon dTAG13 treatment and found that majority of the gene promoters have both factor intensity reduced (Fig. 7f). We identified two subsets of genes with Log2FC in occupancy of both factors <=-1 (MOF sensitive genes) or >=0 (MOF insensitive genes) and confirmed that H4K16ac is diminished at the promoters of most MOF sensitive genes upon MOF depletion, whereas it is retained or gained at most MOF insensitive genes (Fig. 7f,g). GO term analysis revealed that MOF sensitive genes are enriched for the pathways essential for primary cilia function (Fig. 7h), which is consistent with a published study in MOF conditional knockout mice^64^, while MOF insensitive genes are mainly enriched for housekeeping cellular functions (Extended Data Fig. 7d). We also checked whether MOF depletion affected H4K5ac and H4K8ac via ChIP-seq (performed along with ChIP-seq for H4K16ac) in the same cells. Although it appears that only H4K16ac was reduced genome wide (Extended Data Fig. 7e), loci-specific H4K5ac and H4K8ac occupancy was highly correlated with that of H4K16ac; there was little change in H4K5ac and H4K8ac at promoter regions where H4K16ac was retained upon MOF depletion, whereas both H4K5ac and H4K8ac were reduced at promoter regions with diminished H4K16ac (Fig. 7i-k).

**Fig. 7:**
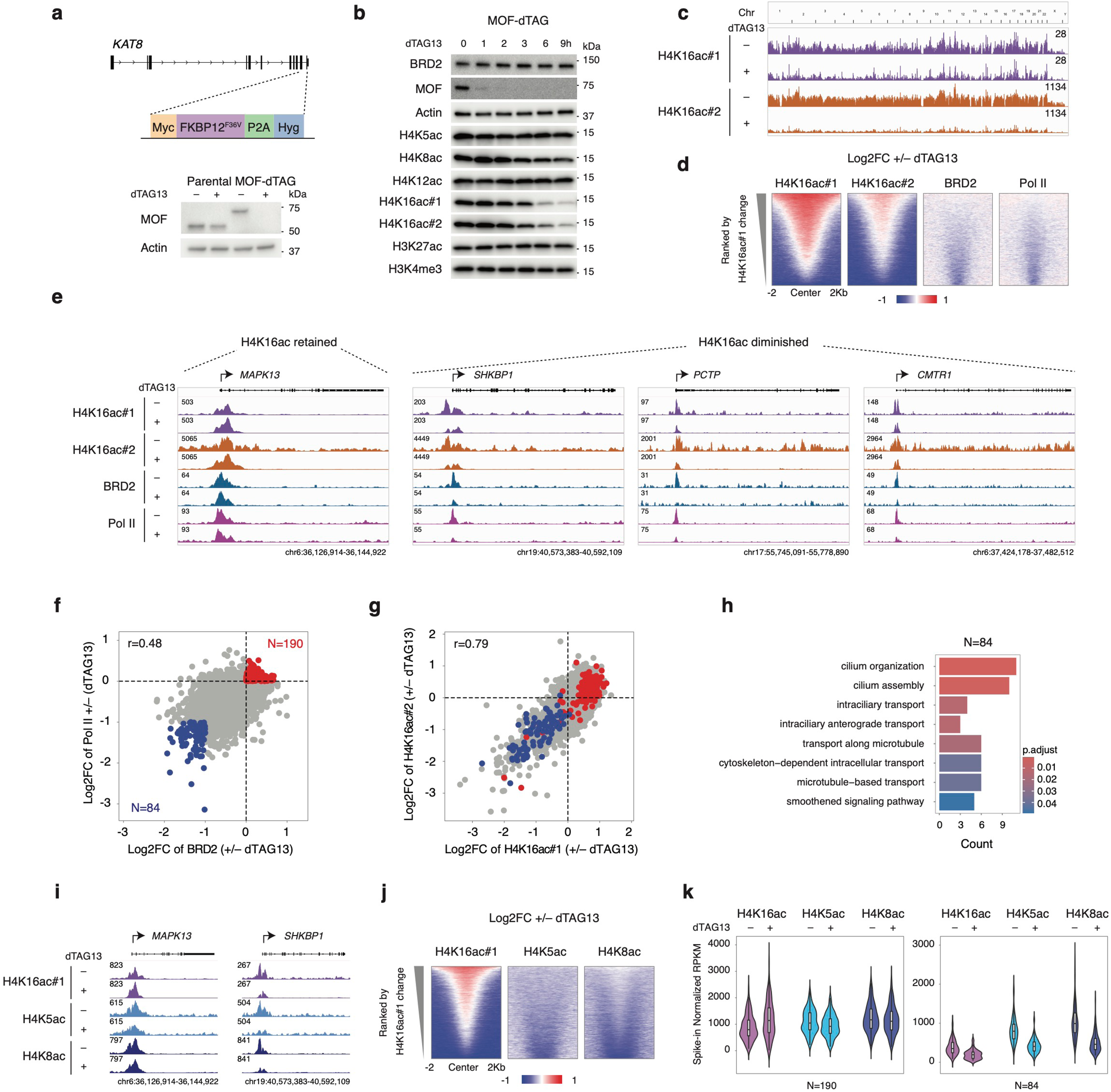
Loss of MOF-mediated H4K16ac leads to decreased BRD2 and Pol II occupancy. (A) Diagram showing the design of MOF-dTAG degron line and western blot confirming MOF depletion upon dTAG13 treatment in the degron line but not parental BRD2-IAA7 cells. Cells were treated with DMSO or 500 nM dTAG13 for 3h. Blots were probed for MOF with actin as loading control. (B) Western blot analysis of dTAG13 treatment time course in the MOF-dTAG line. Cells were treated with 500 nM dTAG13 for 0, 1, 2, 3, 6, or 9h. Blots were probed for BRD2, MOF, H4K5ac, H4K8ac, H4K12ac, H4K16ac (using antibodies from CST (#1) and Abcam (#2)), H3K27ac, and H3K4me3, with actin as loading control. (C) Genomic track visualization of H4K16ac ChIP-seq signals (using antibodies from CST (#1) and Abcam (#2)) across all chromosomes in MOF-dTAG cells treated with DMSO or 500 nM dTAG13 for 9h. (D) Heatmap of log2FC in H4K16ac, BRD2, and Pol II ChIP-seq signals in MOF-dTAG cells treated with DMSO or 500 nM dTAG13 for 9h, at the 23514 shared peak regions (bound by BRD2, Pol II, TBP, and TAF1) identified in Fig. 5b. Regions were ranked by the change in H4K16ac occupancy as detected using antibody #1 (CST). (E) Genomic track visualization of H4K16ac, BRD2, and Pol II ChIP-seq signals upon dTAG13 treatment for 9h at representative gene loci where H4K16ac was retained (*MAPK13*) or diminished (*SHKBP1*, *PCTP*, and *CMTR1*). (F) Scatter plot showing the relationship between the Log2FC in BRD2 occupancy and the Log2FC in Pol II occupancy at the promoters (TSS ± 1kb) of 6481 Pol II transcribed genes upon dTAG13 treatment for 9h in MOF-dTAG cells. Gene promoters with a two-fold or greater reduction in BRD2 and Pol II occupancy are highlighted in blue, while gene promoters that retain occupancy of both factors are highlighted in red. (G) Highlighted gene promoters in panel f were shown in plotting of log2FC of H4K16ac ChIP-seq by two antibodies. (H) GO term pathway enrichment analysis for the 84 genes with 2-fold or greater reduction of BRD2 and Pol II occupancy at promoters, as identified in panel f. (I) Genomic track visualization of H4K16ac, H4K5ac, and H5K8ac ChIP-seq signals at the H4K16ac-retaining (MOF insensitive) gene *MAPK13* and the H4K16ac-diminished (MOF sensitive) gene *SHKBP1* in MOF-dTAG cells treated with DMSO or 500 nM dTAG13 for 9h. (J) Heatmap of log2FC in H4K16ac, H4K5ac, and H4K8ac signal upon 9h dTAG13 treatment for the 23514 shared peak regions identified in Fig. 5b. Regions were ranked by the change in H4K16ac occupancy as detected using antibody #1(CST). (K) ChIP-seq signal intensity for H4K16ac, H4K5ac, and H4K8ac at the promoter regions identified in panel f, in MOF-dTAG cells treated with DMSO or 500 nM dTAG13 for 9h. ChIP-seq signal intensity was calculated as RPKM with spike-in normalization.

**Fig. 8:**
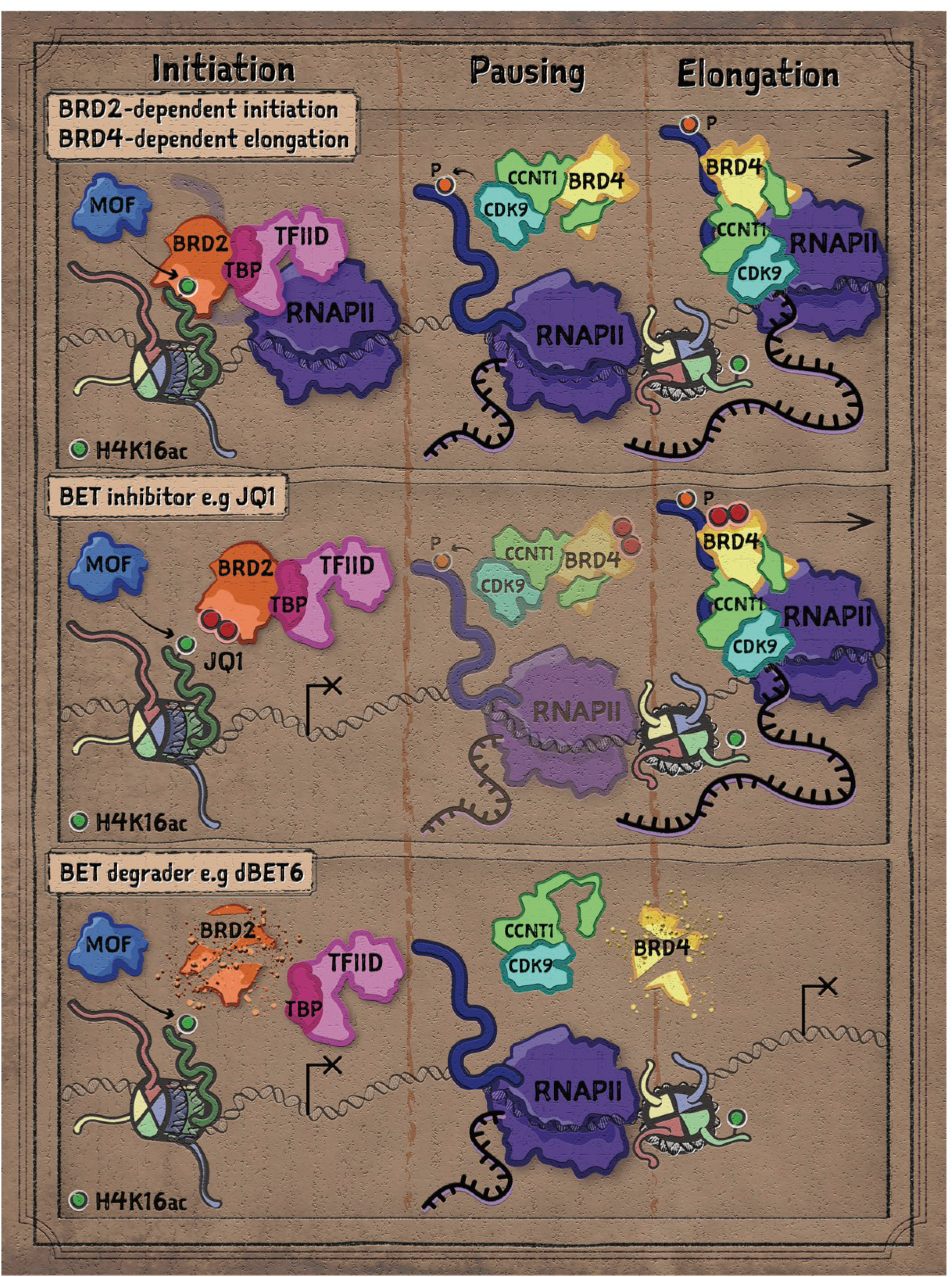
A model of transcriptional regulation by BRD2 and BRD4. We propose distinct but cooperative functions for the BET family proteins BRD2 and BRD4 in the regulation of gene expression in mammalian cells, based on evidence that transcriptional initiation by Pol II is largely dependent on BRD2 but not BRD4, while transcriptional elongation by Pol II is largely dependent on BRD4 but not BRD2. Our results establish a model in which BRD2 bromodomains bind to tetra-acetylated histone H4 harboring the MOF-mediated H4K16ac mark, while the BRD2 C terminus interacts with and recruits TFIID to promote transcriptional initiation by Pol II. Following the post-initiation step in which Pol II pauses at promoter-proximal regions, BRD4 facilitates release of the promoter-proximal paused Pol II by interacting with and enabling the CDK9-containing factor PTEFb to phosphorylate serine residues in the Pol II CTD; we and others have previously established that BRD4 can perform this function without binding to histone acetylation. Small molecule BET bromodomain inhibitors interfere with histone binding and displace BET proteins from chromatin, resulting in a defect in BRD2-mediated transcriptional initiation while BRD4-controlled Pol II release and transcriptional elongation remain intact. Targeted BET protein degraders make use of these same small molecules to target BET proteins, but they create a more profound transcriptional defect because they impact both initiation and elongation by degrading (rather than displacing) both BRD2 and BRD4.

## Discussion

BET proteins are extensively studied factors in biomedical research. BET bromodomain inhibitors are not strongly selective among BET proteins^65^. Despite this fact, and despite the fact that BRD2 is the more abundant BET protein, the mechanism of action for BET inhibitors has been primarily attributed to inhibition of BRD4 bromodomains. However, BRD4’s function in pause release is inherently secondary to establishment of the Pol II machinery in the process of transcription initiation. Our study demonstrates that BRD2 plays an essential role in the regulation of transcriptional initiation at active transcription sites genome wide. Our identification MOF/BRD2/TFIID axis highlights a previously unappreciated role of BRD2 in transcriptional regulation that differs from that of BRD4, providing insight into the mechanism of action for BET inhibitors in both transcriptional initiation and elongation control.

We and others have previously shown in different contexts that the function of BRD4 in Pol II pause-release is largely independent of BRD4 bromodomains^20,66–68^. Our initial findings here suggested that BRD2 bromodomains are largely required for BRD2’s transcriptional initiation-promoting function. Recent studies have demonstrated that the active transcription-associated H3K27ac mark is not required for proper Pol II loading and transcription^69–71^. Because these studies made use of H3K27ac-null cells, in which H4Kac modifications likely remain unaltered, their initially surprising findings could be explained by our finding here, that BRD2 preferentially binds to tetra-acetylated histone H4 harboring H4K16ac (Fig. 6). However, it remains intriguing that even when MOF is depleted to the point that it is undetectable by Western blot analysis and genome wide reduction of H4K16ac is observed, many transcription loci retained or even gained H4K16ac (Fig. 7). Future screening studies are needed to determine if another HAT such as TIP60^72^ (or some novel HAT) compensates for the loss of MOF, or whether other factor(s) might block the accessibility of H4K16ac loci to HDACs such as those in the sirtuin family^73,74^. Among H4Kac, our impression tends to favor H4K16ac as the major determinant of BRD2 chromatin binding given that H4K16ac is the most highly correlated with BRD2 occupancy (Fig. 6h), and that H4K5ac and H4K8ac appear to be slightly diminished at promoters where BRD2 and Pol II occupancy is retained upon MOF depletion. However, more detailed future studies are required to test this hypothesis with specific depletion of non-catalytic subunits in the MOF complexes NSL or MSL.

Our data suggests that BRD3 can apparently play a similar role to that of BRD2, as its overexpression is able to compensate for the loss of BRD2 to facilitate transcription initiation in DLD-1 cells (Fig. 6). The likely redundancy between BRD2 and BRD3 is also supported by a positive correlation of CRISPR score for BRD2 with the expression level of BRD3, but not BRD4 (Extended Data Fig. 7h). However, we were able to identify BRD2 function via BRD2 depletion alone because endogenous BRD3 expression levels are much lower than BRD2 in DLD-1 cells (Extended Data Fig. 2a and Fig. 7f). This difference in expression level appears to be a general phenomenon, because BRD2 expression is higher than that of BRD3 (median ∼10X than BRD3) in all 1103 cell lines available in the DepMap database (Extended Data Fig. 7f,g).

The fact that the yeast BET homologues Bdf1/2 facilitate the recruitment of TFIID indicates that BRD2/3’s role in transcription initiation is a conserved function^34^. We hypothesize that during evolution, gain of the PTEFb interacting domain in the C terminal of BRD4 conferred a novel function in Pol II elongation control that led to eventual loss of the initiation function preserved in BRD2/3. We identified the C terminus of BRD2 as required for interaction with TFIID and critical to BRD2’s initiation-promoting function. Interestingly, the BRD2 C terminus region contains a predicted coiled-coil domain that is conserved from yeast to human and was previously indicated by Werner et al.^58^ to be essential for BRD2’s growth-promoting function in murine erythroblasts, although the nature of this function was not known. Our study provides a sound explanation for why the BRD2 C terminal domain was required for growth in these cells.

Our ChIP-seq studies suggest that Pol II chromatin occupancy appears to be highly correlated with the level of BRD2 occupancy at the same sites. For example, GFP-BRD2FL overexpression levels are higher in BRD2-AID cells than in BRD2-IAA7 DLD-1 cells (Fig. 3a and Fig. 4g). Consequently, Pol II occupancy under the rescue condition, where endogenous BRD2 is replaced by GFP-BRD2FL, is higher in the AID than in the IAA7 degron line (Fig. 3c and Fig. 4i). Pol II ChIP-seq upon JQ1 treatment at 1 uM in MOLT4 cells vs 10 uM in DLD-1 cells also revealed the disparate degree of Pol II loss (Fig. 1c,d and Fig. 5). This finding has important implications for physiology or therapeutic application, as BET protein levels can vary in different tissues or conditions^31,63^. In prostate cancer, mutations in the E3 ligase component SPOP can stabilize BET proteins (which are otherwise targeted for degradation via an intrinsic degron sequence), conferring resistance to treatment with BET inhibitor^75,76^. Phosphocreatine binds to and protects BRD2 from SPOP mediated degradation, and disruption of phosphocreatine biosynthesis sensitizes glioblastoma to BET inhibitor treatment^32,58^. In both cases, we speculate that the level of BRD2/3 occupancy could be a determinant of resistance versus sensitivity to BET inhibition.

NUT carcinoma (NC) is an aggressive squamous carcinoma characterized by the fusion of *NUT* gene with another gene such as BRD4. The BRD4-NUT oncofusion protein, which drives a particularly aggressive form of midline carcinoma, establishes massive regions of histone hyperacetylation called mega domains via interaction with the histone acetyltransferase p300^77–80^. Patients with non-*BRD4::NUTM1* (e.g. *BRD3::NUTM1* and *NSD3::NUTM1*) tumors generally have a better prognosis and respond better to BET inhibitor treatment than patients with BRD4-NUT positive tumors^81,82^. Consistent with this, a novel case of BRD2-NUT fusion carcinoma was recently reported to be less aggressive and to respond to chemotherapy exceptionally well^83^. Critically, the BRD2-NUT fusion in this specific case affects the BRD2 C terminus, which we demonstrate here to contain the TFIID interacting domain (Fig. 4). Considering the distinct function of BRD2/3 vs BRD4 in transcription regulation, it is encouraging to speculate that by interfering with transcription initiation, loss of TFIID binding capacity in this specific BRD2-NUT fusion could affect transcriptional capacity, leading to a less aggressive condition. It would be necessary to elucidate how BRD2-NUT fusion affects the interaction between BRD2 and TFIID, including work to map the specific BRD2 C terminal residues required for TFIID binding; defining the BRD2-TFIID interaction surface could potentially enable novel therapeutic development for diseases such as NUT carcinoma. Given the preferential binding of BRD2 bromodomains to tetra-acetylated histone H4, future studies should also examine the impact of H4Kac and other histone acetylation profiles apart from H3K27ac.

## Materials and Methods

### Tissue culture

DLD-1 (CCL-221) human cells were purchased from ATCC. BRD2-AID and BRD4-IAA7 DLD-1 cells were from previous studies^19,20^. BRD2-IAA7 and MOF-dTAG DLD-1 degron cells were generated as described in the following section. All cell lines were cultured in DMEM (Corning, #10013CV), supplemented with 10% FBS (Sigma-Aldrich, #F2442), 1% Glutmax (Gibco, #35050061), and 1% PS (Gibco, #15140122), in 37°C incubator with 5% CO2.

### Generation of degron cell lines

BRD2-IAA7 degron differs from BRD2-AID degron by updating the degron peptide and their auxin receptor F-box (AFB) protein from mini-AID and OsTIR1 to mini-IAA7 and AtAFB2^18^. BRD2-IAA7 degron cells were generated similarly to previously described^20^. Briefly, PX330 (Addgene, #42230) with the insertion of sgRNA (TGATTCAGACTCAGGCTAAG) targeting the stop codon area of the BRD2 genomic locus was co-transfected with a donor plasmid using the Lipofectamine 3000 Transfection Reagent (Invitrogen, #L3000001) to trigger donor integration at the target site via homologous recombination repair. The donor plasmid, which we designed to integrate the IAA7 tag after the final BRD2 exon and introduce the F-box protein AtAFB2 under the EF1a promoter, was made using the pBlueScript II SK (+) backbone and synthesized gBlocks for the IAA7-AtAFB2 pair and antibiotic selection marker. Upon transfection, DLD-1 cells were selected for colony formation in the presence of Geneticin (Gibco, #10131027) for 2 weeks. Single clone colonies were picked, and genomic DNA was extracted using Zymo kit (#D3011). Homozygous knock-in was verified by PCR and western blot analysis. MOF-dTAG degron cells were generated similarly with donor plasmid synthesized by Twist Biosciences harboring the homology arms and the Myc-FKBP12^F36V^-P2A-Hygromycin sequences in between. PX330 with insertion of sgRNA (TCTCCAAGAAGTGAGCAGCC) was co-transfected with the donor plasmid in BRD2-IAA7 DLD-1 cells. Hygromycin B (Gibo, #10687010) was used for the selection of knock-in clones.

### Drug treatments

Auxin (#ab146403) was purchased from abcam. Doxycycline (#72742) was obtained from Stem Cell Technologies. dEBT6 (#S8762) was purchased from Selleckchem. JQ1 (#4499) and dTAG13 (#6605) were purchased from Tocris.

### Generation and expression of BET and mutant constructs

TetOn lentiviral vector (#110280) and BRD2-FL cDNA (#65376) were obtained from Addgene. The BRD2-ΔBD mutant was generated from the BRD2-FL construct using the Q5 Site-Directed Mutagenesis Kit (NEB, #E0554S). BRD4-FL and BRD4-ΔBD constructs were from previous study^20^. All other BET and mutant constructs were generated via NEBuilder HiFi DNA assembly reaction (NEB, #E2621S) with synthesized gBlocks from Twist Biosciences. BET and mutant lentiviral constructs were amplified by transformation of stable competent E. coli (NEB, #C3040H). All construct plasmid insertions were verified by Sanger sequencing (ACGT). Lentivirus for transducing mammalian expression was achieved by co-transfecting the BET and mutant expressing plasmids with pspax2/pmd2.g lentiviral packaging plasmids in 293T cells in 6-well format. Lentivirus was collected and filtered for infection in 6-well plates. For construct expression in BRD2-AID DLD-1 cells, due to the resistance to common selective markers, single clones were picked using GFP signals upon Dox induction. For construct expression in BRD2-IAA7 DLD-1 cells, infected cells were selected with Blasticidin (Gibco, #A1113903) for over two weeks. BET and mutant expression were achieved by adding 50 nM Dox into the medium and incubating for 36 or 48h.

### MOF knockdown by CRISPR

LentiCRISPR v2 vector (Addgene, #52961) was modified by inserting the sgRNA sequence (CCATACTTACAGTTAACCGG). Lentiviral packaging and infection were as described above. Infected BRD2-IAA7 DLD-1 cells were selected with Puromycin Dihydrochloride (Gibco, #A1113803) for over three days.

#### Western blot

Whole cell lysates for western blot were prepared by directly lysing the cells with Laemmli sample buffer (Bio-Rad, #1610747) and boiling for 10 minutes. BRD2 (#5848), TBP (#44059), TAF1 (#12781), BRD4 (#13440), BRD3 (#94032), β-Actin (#3700), V5-Tag (#13202), CDK9 (#2316), Cyclin T1 (#81464), KAT8 (#46862), H4K5ac (#8647), H4K8ac (#2594), H4K12ac (#13944), H4K16ac#1 (#13534),, H3K27ac (#8173), and H3K4me3 (#9751) antibodies were purchased from Cell Signaling Technology. TAF6 (#A301-276A) antibody was obtained from Fortis Life Sciences. H4K16ac#2 (#ab109463) was purchased from Abcam. GFP Antibody (#sc-9996) was purchased from Santa Cruz Biotechnology.

### In vitro recombinant nucleosome binding assay

GST-tagged bromodomains of BRD2 were purified by GenScript. Sequences for the domains are as follows:

GST-BD1:

MSPILGYWKIKGLVQPTRLLLEYLEEKYEEHLYERDEGDKWRNKKFELGLEFPNLPYYI DGDVKLTQSMAIIRYIADKHNMLGGCPKERAEISMLEGAVLDIRYGVSRIAYSKDFETLK VDFLSKLPEMLKMFEDRLCHKTYLNGDHVTHPDFMLYDALDVVLYMDPMCLDAFPKLVCFKKRIEAIPQIDKYLKSSKYIAWPLQGWQATFGGGDHPPKSDRVTNQLQYLHKVVMKALWKHQFAWPFRQPVDAVKLGLPDYHKIIKQPMDMGTIKRRLENNYYWAASECMQDFNTMFTNCYIYNKPTDDIVLMAQTLEKIFLQKVASMPQEE

GST-BD2:

MSPILGYWKIKGLVQPTRLLLEYLEEKYEEHLYERDEGDKWRNKKFELGLEFPNLPYYI DGDVKLTQSMAIIRYIADKHNMLGGCPKERAEISMLEGAVLDIRYGVSRIAYSKDFETLK VDFLSKLPEMLKMFEDRLCHKTYLNGDHVTHPDFMLYDALDVVLYMDPMCLDAFPKLVCFKKRIEAIPQIDKYLKSSKYIAWPLQGWQATFGGGDHPPKSDKLSEQLKHCNGILKELLSKKHAAYAWPFYKPVDASALGLHDYHDIIKHPMDLSTVKRKMENRDYRDAQEFAADVRLMFSNCYKYNPPDHDVVAMARKLQDVFEFRYAKMPDE

Nucleosome binding assays were carried out by EpiCypher.

#### Fluorescent imaging and immunofluorescence staining

Upon induction of GFP-tagged constructs, cells in the cell culture dish were either directly imaged or fixed by 4% Formaldehyde (ThermoFisher Scientific, # 50-980-495) in PBS at room temperature with shaking for 20 minutes before imaging. For immunofluorescence staining, cells cultured in chamber wells (ThermoFisher Scientific, #154526) were washed with PBS twice and fixed by 4% Formaldehyde for 20 minutes. Fixed cells were then washed 1x with PBS and incubated with 0.2% Triton X-100 in PBS at room temperature for 10 minutes with shaking. 10% Normal Goat Serum blocking solution (Vector Laboratories, #S-1000-20) in 0.1% Tween 20 was used for 30-minute blocking. To stain BRD2 foci, cells were incubated with Alexa Fluor 488 conjugated Anti-BRD2 antibody (Abcam, #ab197865) in blocking solution over night at 4°C and mounted with Antifade Mountant with DAPI (Invitrogen, # P36962). Cells then were washed with PBS for 3x 5 minutes before mounting and imaging. Nikon DS-Qi2 and DS-Fi3 cameras were used for imaging. For overexpressed constructs, since the expression level varies and the signal intensity was not compared, exposure time was determined automatically. For BRD2 IF antibody validation, exposure time was set at 80ms for –/+dBET6 treatment in NCIH2009 cells.

#### GFP IP and IP-MS

Similar to previous study^20^, to prepare the GFP IP samples, cells were trypsinized and washed with PBS twice before lysed in cold Triton X-100 lysis buffer (50 mM Tris HCl, pH 8.0, 150 mM NaCl, 1.5 mM MgCl2, 10% Glycerol, 0.5% Triton X-100, DTT (ThermoFisher Scientific, #A39255), Benzonase (Millipore Sigma, #E1014), Protease inhibitor (ThermoFisher Scientific, #PIA32963), and Phosphatase inhibitor (ThermoFisher Scientific, #PIA32957)) followed by rotation for 45min and centrifugation at 20,000g for 15 minutes at 4°C. The supernatant was incubated with ChromoTek GFP-Trap magnetic beads (Proteintech, #gtd or #gtma) for > 4h at 4°C for immunoprecipitation, then beads were washed 5x with cold Triton X-100 lysis buffer for IP western, or 4x with cold Triton X-100 lysis buffer plus 1x with freshly prepared cold 100mM NH4HCO3 (Millipore Sigma, #A6141) for IP MS. For IP western, 2X Laemmli sample buffer was directly added to the washed beads, followed by boiling for 10 minutes. For IP-MS, 2.5 M Glycine pH 2.0 was used to elute the immunoprecipitated proteins and Tris buffer pH>10 was used to neutralize the eluate at RT. Neutralized eluate was snap frozen for MS or mixed with Laemmli sample buffer and boiled for 5 minutes for western blot. MS was carried out by the Proteomics Core Facility at the University of Arkansas for Medical Sciences.

#### ChIP-seq

ChIP-seq experiments were carried out similarly as previously described^20^. Briefly, cells in a 15cm plate were crosslinked by 15 ml 1% PFA (ThermoFisher, #28908) in PBS with shaking for 10 minutes at room temperature. 2 ml of 2.5 M Gylcine was added to quench the reaction by shaking for 5 minutes. Crosslinked cells were then washed with PBS twice before scraping and collecting. Covaris E200 was used for chromatin sonication with following settings: 10% duty factor, 200 cycles per burst, and 140 peak intensity power, 10 minutes. Antibodies used for ChIP are as follows: Pol II (Rpb1 NTD) (Cell Signaling Technology, #14958), H3K27ac (Cell Signaling Technology, #8173), BRD4 (Fortis Life Sciences, #A301-985A50), GFP (DSHB, cocktail of #GFP-G1, #GFP-12A6, #GFP-12E6, and #GFP-8H11 or Cell Signaling Technology, #55494), BRD2 (Cell Signaling Technology, #5848), TBP (Cell Signaling Technology, #44059), TAF1 (Cell Signaling Technology, #12781), H4K5ac (Cell Signaling Technology, #8647), H4K8ac (Cell Signaling Technology, #2594), H4K12ac (Cell Signaling Technology, #13944), H4K16ac#1 (Cell Signaling Technology, #13534), H4K16ac#2 (Abcam, #ab109463). Spike-in chromatin (Active Motif, #53083) and antibody (Active Motif, #61686) for normalization were added before immunoprecipitation. Alternatively, chromatin isolated from mouse embryonic fibroblast cells was used. For immunoprecipitation, antibodies were incubated with the lysate over night at 4°C with rotation for immunoprecipitation. Protein G-coupled Dynabeads (Invitrogen, #10004D) were used to pull down the immune complexes. Protease K (Roche, #3115828001) was used during reverse crosslink (65°C, 1200 rpm, overnight). DNA was then purified with either QIAquick PCR purification kit (Qiagen, #28106) or ChIP DNA Clean & Concentrator kit (Zymo Research, #D5205). Libraries were prepared by using the KAPA HTP library preparation kit (Roche, #07961901001) and sequenced on the Illumina NovaSeq 6000 or NextSeq 2000.

### RNA-seq

Similarly to previously described^67^, DLD-1 cells were cultured in 6 well plate and total RNAs were extracted using the RNeasy Mini Kit (QIAGEN, #74106) according to the procedure. mRNAs were enriched using the NEBNext poly(A) mRNA magnetic isolation module (NEB, #E7490L). Libraries were prepared using the NEBNext Ultra II Directional RNA library preparation kit (NEB, #E7760L) and sequenced on NextSeq 2000. Quality trimming was done using Trimmomatic^84^ with parameters TRAILING:30 MINLEN:20. Reads were aligned to the hg38 genome using STAR v.2.5.2^85^, and only uniquely mapped reads with a two-mismatch threshold were kept. Output BAM files were converted into bigWig track files to display coverage throughout the genome (in rpm).

#### PRO-seq

As previously described^86,87^, PRO-seq experiments for BRD4 depletion and dBET6 treatment were performed according to the published original and modified protocols respectively^88,89^. Briefly, DLD-1 cell nuclei (10 million for BRD4 depletion and 1 million for dBET6 treatment) were mixed with Drosophila S2 spike-in nuclei, which were then used for nuclear run-on assay and subsequent library preparation. Libraries were sequenced on Illumina NextSeq 500 or NovaSeq 6000. Low-quality bases and adapters from 3’ ends were removed using cutadapt 1.14^90^ requiring a read length of 16–36 bp. Ribosomal RNA reads were also removed. The remaining reads were aligned using bowtie 2.2.6 with --very-sensitive option^91^ to a concatenated genome comprised of human hg38 and fly dm6 assemblies. 5’ ends of aligned reads with MAPQ ≥ 30 were retained and assigned to the opposite strand. Read counts were normalized to total reads aligned to the spike-in genome for the output of bigwig files.

#### TT-seq

Similar to previously described^92^, ∼18 million cells after Dox treatment for 48h for expression of GFP or GFP-tagged BRD2 FL and ΔBD mutant were treated with or without Aux for 3h. Nascent RNA labeling was achieved by adding 500 μM 4-thiouridin (4sU) (Sigma, #T4509) to the media and incubating for 10 minutes. TT-seq library was prepared according to published studies^77,93^. Briefly, following 4sU labeling, total RNA was extracted using TRIzol reagent (ThermoFisher, #15596026). Spike-in RNA prepared the same way from Drosophila S2 was added. Mixed RNA was fragmented using Magnesium RNA Fragmentation Module (NEB, #E6150S), followed by biotinylation of 4sU using Biotin-XX-MTSEA (Biotium, #90066) in 20% DMF. Biotinylated RNA was enriched using Dynabeads MyOne Streptavidin C1 (ThermoFisher, #65001), followed by reduction of disulfide bonds in 100 mM DTT for RNA elution. DNA libraries were prepared from the enriched RNA fragments and sequenced on NextSeq 2000. Low-quality bases were removed from 3′ ends using cutadapt 1.14^90^. Reads were aligned to concatenated genome of human hg38 and fly dm6 with Ensembl annotation data release 103 using STAR 2.7.5^85^. Read counts were normalized to total reads aligned to the spike-in genome for the output of bigwig files.

### Data Analysis

#### ChIP-seq

ChIP-seq reads were aligned to hg38 and dm6 with bowtie 1/2^94^. BamCoverage in deeptools 3.1.1^95^ was used to extend ChIP-seq reads to 150bp. Normalization was carried out using fly spike-in reads except for Pol II ChIP-seq in NCIH2009 (Fig. 1c,d) and BRD4 rescue experiment (Extended Data Fig. 3b) which were carried out using mouse spike-in reads. GFP ChIP-seq was normalized to total mapped reads except for JQ1 treatment in Fig. 2d,f. Spike-in normalization was not used for BRD2, H3K27ac, and histone H4 acetylation ChIP-seq in Fig. 6g,h. All other ChIP-seq were normalized to spike-in. Log2FC of ChIP-seq was calculated using bigwigCompare in deeptools with the following nondefault options: binSize 10, pseudocount 0.1. Heatmaps and metagene plots were also generated in deeptools. Genes (N=6,481) with TSS and TES annotation were obtained from our previously published study^96^. BET-bound enhancers (N=8,149) previously isolated from hg19 alignment were converted to hg38 by using the UCSC Genomic“Lift Genome Annotations” online tool. FeatureCounts 2.0.1^97^ was used to calculate the mapped read counts at given loci. The output read counts were then normalized to total mapped reads of spike-in and the region length to get the binding density. Pol II reduction percentage was calculated using Pol II binding density as follows: (Control - Treatment)/Control *100%. Mean occupancy density was the average binding density of regions with a specific Pol II reduction value. Peak calling was performed using MACS2/2.1.0^98^ with a threshold of q=0.01. Peak annotation was carried out with homer/5.1^99^. Intervene 0.6.4^100^ was used to generate the Venn diagram and shared peak regions. To isolated Pol II transcribed genes in MOLT4 and NCIH2009 cells, called Pol II ChIP-seq peaks were overlapped with annotation from GENCODE release 43^101^. Lowly expressed genes with RNA-seq TPM <10 (for MOLT4) or htseq count <10 (for NCIH2009) were filtered out. All tracks were visualized in igv 2.13.2 (Broad Institute).

#### PRO-seq

Heatmaps for PRO-seq were generated in deeptools similarly to ChIP-seq with only minor difference: bigwigCompare for Log2FC of PRO-seq signals and computeMatrix for heatmap were calculated separately for plus and minus strand. The output matrices were then merged using computeMatrixOperations rbind function within deeptools.

#### TT-seq

TT-seq read counts within the genebody from TSS to TES were also calculated using FeatureCounts 2.0.1 with following setting: -p -B -C -s 2. The output read counts were then normalized by total mapped spike-in reads. The MA and Gene Ontology plots were generated in R. Mild cutoff for up and down regulated gene selection was set at |log2FC| of 0.415. Among the down regulated genes, BD sensitive genes were further selected as log2FC of (FL_auxin vs GFP_auxin) >= 0.415 and log2FC of (ΔBD_auxin vs FL_auxin) <= −0.415. BD insensitive genes were selected as log2FC of (FL_auxin vs GFP_auxin) >= 0.415 and log2FC of (ΔBD_auxin vs FL_auxin) >=0.

## Acknowledgement

The authors thank Brianna Monroe for illustrating the model. We are grateful to all the members in the Shilatifard for the insightful discussion for the study. We thank Yue Feng, Ruli Gao, and Daniel Foltz for providing equipment and reagents. Studies in the Shilatifard lab are supported by generous funding through Chan Zuckerberg BioHub in Chicago and the Outstanding Investigator Award mechanism by NCI grant R35-CA197569.

## Author contributions

B.Z. and A.S. conceived and designed the experiments. B.Z. and R.Q. carried out the experiments. B.Z. performed data analysis and bioinformatics. M.I., B.H., and M.D. performed NGS. Y.A. helped with the PRO-seq and TT-seq experiments. B.Z. wrote and S.G. revised the manuscript. B.Z., S.G., and A.S. finalized the manuscript.

## Competing interests

All authors declare that they have no competing interests.

---

**Extended Data Fig. 1:**

(A) Metagene plots of average PRO-seq signal at Pol II-transcribed gene promoters (n=6481) and at BET-bound enhancers (n=8147, as determined in previous work^19^; see Methods), in BRD4-AID DLD-1 cells treated with H2O or auxin (Aux, 500 uM) for 3h and in WT DLD-1 cells treated with DMSO or dBET6 (250 nM) for 2h.

(B) Heatmaps of Pol II occupancy at 7912 Pol II-transcribed genes (ranked by Pol II occupancy) in NICH2009 cells that were treated with DMSO or with dBET6, with or without complementation (rescue) via expression of the GFP-BRD4ΔBD construct. Heatmaps of log2-transformed fold change (Log2FC) in ChIP-seq signal between conditions are shown at right. Pol II ChIP-seq in NCIH2009 cells was previously published^20^, but realigned and normalized to spike-in control.

**Extended Data Fig. 2:**

(A) Genomic track visualization of mRNA-seq signal (two replicates) on either strand at the *BRD2*, *BRD3*, and *BRD4* gene loci in DLD-1 cells, indicating the relative expression levels of these BET family proteins.

(B) Immunofluorescent staining for endogenous BRD2 in NCIH2009 cells treated with DMSO or 250 nM dBET6 for 3h. Scale bar, 50 um.

(C) Immunofluorescent staining for endogenous BRD2 in NCIH2009 cells treated with DMSO or 10 uM JQ1 for 24h. Scale bar, 50 um.

(D) Immunofluorescent staining for endogenous BRD2 in 293T cells treated with DMSO or JQ1 (1 uM, 5 uM, 10 uM) for 3h. Scale bar, 50 um.

(E) Genomic track visualizations of GFP ChIP-seq signal (BRD4FL or BRD4ΔBD occupancy) at the representative promoter (*MAPK13*) and enhancer in BRD4-IAA7 DLD-1 cells expressing the TetOn-NLS-GFP vector or the GFP-BRD4FL or GFP-BRD4ΔBD constructs. Heatmaps of BRD4FL or BRD4ΔBD occupancy and of Log2FC compared to the vector control at Pol II-transcribed gene promoters (n=6481) and at BET-bound enhancers (n=8147).

**Extended Data Fig. 3:**

(A) Heatmaps of Pol II ChIP-seq signal and Log2FC for the BRD2 depletion and rescue as well as JQ1 treatment and washout at enhancers.

(B) Heatmaps of genome-wide Pol II ChIP-seq signal and Log2FC at both promoters and enhancers for BRD4 depletion in the BRD4-IAA7 DLD-1 degron line and rescue with BRD4FL or BRD4ΔBD.

(C) MA plot for genome-wide differential transcription in the auxin-treated vs untreated condition, from the TT-seq described in Fig. 3d. On the y-axis, M is the log2FC of normalized TT-seq read counts within gene bodies (from TSS to TES) with vs without auxin. On the x-axis, A is the average of log2 transformed normalized TT-seq read counts for the treated and untreated conditions. Upregulated genes (N=111) with log2FC ≥ 0.415 are highlighted in red and downregulated genes (BRD2-dependent genes, N=1883) with log2FC ≤ −0.415 are highlighted in blue. *CYP1A1* is annotated as a known auxin-induced gene. *MAPK13* and *CYLD* (signal tracks shown in Fig. 3d) are also annotated.

(D) Metagene plot of TT-seq signal corresponding to both sense and antisense transcription at BD sensitive and insensitive BRD2-dependent genes (as identified in Fig. 3f), for each of the depletion and rescue conditions described in Fig. 2d. (Note that antisense transcription for *PARP2* was excluded from metagene plot due to the abnormally high reads for *RPPH1* located upstream of *PARP2*).

(E) Metagene plot and heatmap of log2FC in Pol II ChIP-seq signal upon rescue with BRD2ΔBD vs BRD2FL, for the BD-sensitive and BD-insensitive BRD2-dependent genes identified in Fig. 3f.

(F) GO term enrichment analysis of the 450 BD-sensitive genes identified in Fig. 3f.

**Extended Data Fig. 4:**

(A) Western blot analysis of GFP-IP, performed using one of the six sets of GFP IP-MS sample replicates in Fig. 4a. Blots were probed for BRD2 and the TFIID subunit TBP.

(B) Ranked plot of the top 100 genes for which CRISPR effect score is most positively correlated with that of *TAF5*, *TAF8*, or *TAF3* across all cell lines available from DepMap. Genes encoding TFIID subunits are named and highlighted in red; *BRD2* is highlighted in blue.

(C) Schematic illustration of the BRD2FL construct and additional deletion mutant constructs expressed from the TetOn-NLS-GFP vector, with western blot analysis of construct expression levels in WT DLD-1 cells. ΔBDET: BRD2 with both BD and ET domain deletion. C1: ΔBDET with additional deletion eliminating regions flanking bromodomain 1 (BD1). C2: ΔBDET with additional deletion eliminating all bromodomain-flanking regions. Western blot was performed using both BRD2 (top) and GFP (bottom) antibodies because the FL band was difficult to detect via the GFP antibody.

(D) Western blot analysis of GFP IP from WT DLD-1 cells expressing the GFP-tagged BRD2FL, BRD2ΔBDET, BRD2C1, or BRD2C2 constructs. Blots were probed for the TFIID subunits TAF1, TAF6, and TBP.

(E) Genomic track visualization of BRD2 and Pol II ChIP-seq signals at the *MAPK13* promoter in BRD2-IAA7 DLD-1 cells treated with H2O or 500 uM auxin for 3h.

(F) Heatmap of BRD2 and Pol II ChIP-seq signal and Log2FC in signal upon BRD2 depletion at the promoters of 6481 Pol II-transcribed genes in BRD2-IAA7 DLD-1 cells treated with H2O or 500 uM auxin for 3h.

(G) Scatter plot comparing the percentage reduction in Pol II ChIP-seq signal between the BRD2-IAA7 and BRD2-AID degron lines treated with 500 uM auxin for 3h, at the 6466 gene promoters where Pol II signal was reduced in both lines.

(H) Genomic track visualization of GFP, Pol II, TBP, and TAF1 ChIP-seq signals at three distal enhancers in untreated and auxin-treated BRD2-IAA7 DLD-1 cells expressing the TetOn-NLS-GFP vector and in auxin-treated BRD2-IAA7 DLD-1 cells expressing the GFP-tagged BRD2 FL or ΔC constructs.

(I) Heatmaps of GFP, Pol II, TBP, and TAF1 ChIP-seq signal at BET-bound enhancers (n=8147), with heatmaps of log2FC in signal upon endogenous BRD2 depletion and upon rescue with the GFP-tagged BRD2 FL or ΔC constructs, for the conditions described in panel h.

**Extended Data Fig. 5:**

(A) Genomic track visualization of BRD2, Pol II, TBP, and TAF1 ChIP-seq signals at a distal enhancer region in DLD-1 cells treated with DMSO or 10 uM JQ1 for 3h.

(B) Heatmaps of ChIP-seq signal for BRD2, Pol II, TBP, and TAF1 at Pol II-transcribed gene promoters (n=6481) and BET-bound enhancers (n=8147) in DLD-1 cells treated with DMSO or 10 uM JQ1 for 3h, with heatmaps of log2FC in signal upon JQ1 treatment shown for each factor.

(C) Mean occupancy density plot for BRD2, Pol II, TBP, and TAF1 ChIP-seq signals at Pol II-transcribed gene promoters and BET-bound enhancers in DLD-1 cells treated with DMSO or 10 uM JQ1 for 3h, ordered by their percentage reduction in Pol II ChIP-seq signal. ChIP-seq signals were normalized to spike in control and peak region length, and log2 transformed.

(D) Scatter plot showing the correlation between Log2FC in BRD2 ChIP-seq signal and the Log2FC in Pol II, TBP, and TAF1 ChIP-seq signals upon JQ1 treatment for the 67305 peak regions unique to Pol II, as shown in Fig. 5b (55011 were plotted after removing NA values). Pearson scores are shown for each correlation.

(E) Metagene plots and heatmaps of log2FC in Pol II ChIP-seq signal upon 10 uM JQ1 treatment in DLD-1 cells, for the BD-sensitive and BD-insensitive BRD2-dependent genes identified in Fig. 3f.

**Extended Data Fig. 6:**

(A) Western blot of GFP IP from DLD-1 cells expressing GFP-tagged BRD2, BRD3, or BRD4 constructs (all full-length). Blots were probed for the TFIID subunits TAF1, TAF6, and TBP.

(B) Fluorescence microscopy images of GFP signal in BRD2-IAA7 DLD-1 cells following the Dox-induced expression of the GFP-tagged BET proteins and mutant constructs illustrated in Fig. 6a. Scale bar, 10uM.

(C) Genomic track visualization of Pol II ChIP-seq signal at BET protein gene loci in BRD2-IAA7 DLD-1 cells, for the BRD2 depletion and rescue conditions described in Fig. 6c.

(D) Genomic track visualization of GFP ChIP-seq signal at representative gene loci (*MAPK13* and *CSF1*) in auxin-treated BRD2-IAA7 DLD-1 cells expressing GFP-tagged BRD2 or the GFP-tagged bromodomain swap constructs (BRD4x2BD or BRD2x4BD).

(E) Heatmaps of GFP ChIP-seq signal at the promoters of 6481 Pol II-transcribed genes for the conditions in panel d.

(F) Genomic track visualization of GFP and BRD4 ChIP-seq signals at representative gene loci (*MAPK13* and *CSF1*) and distal enhancers upon BRD2 depletion by auxin treatment in BRD2-AID cells.

(G) Heatmaps of GFP and BRD4 ChIP-seq signals at 6481 Pol II-transcribed gene promoters (n=6481) and 8147 BET-bound enhancers (combined into single plots) upon BRD2 depletion by auxin treatment in BRD2-AID cells, with heatmaps of log2FC in signal for auxin-treated vs untreated cells.

(H) Oriole staining for the purified GST-tagged BD1 and BD2 domains of BRD2.

**Extended Data Fig. 7:**

(A) Western blot analysis of MOF knockdown by CRISPR in untreated sgCtrl versus sgKAT8 BRD2-IAA7 DLD-1 cells, and the impact of MOF knockdown on the tetra-acetylation of H4. Blots were probed for MOF, H4K5ac, H4K8ac, H4K12ac, and H4K16ac, with actin as loading control.

(B) Genomic track visualization of BRD2 and Pol II ChIP-seq signal at the *SHKBP1* gene locus in sgKAT8 compared to sgCtrl cells.

(C) Heatmap of log2FC in BRD2 and Pol II ChIP-seq signals upon MOF knockdown (sgKAT8 vs sgCtrl) at the 23514 shared peak regions bound by BRD2, Pol II, TBP, and TAF1, as identified in Fig. 5b.

(D) GO term pathway enrichment for the 190 genes with promoters that retained BRD2 and Pol II occupancy following 9h dTAG13 treatment in MOF-dTAG cells (as identified in Fig. 7f).

(E) Genomic track visualization of H4K16ac (as detected by antibody #1), H4K5ac, and H4K8ac ChIP-seq signals across all chromosomes in MOF-dTAG cells treated with DMSO or dTAG13 for 9h.

(F) Scatter plots showing the relationship between the log2-transformed expression level (TPM+1) of BRD2 and that of BRD3 (left) or BRD4 (right), across a total of 1103 cell lines available from DepMap. Red dashed lines corresponding to equal expression levels are shown for comparison. The DLD-1 cell line is also highlighted in red.

(G) Violin plot of Log2FC in the expression level (TPM+1) of BRD2 vs that of BRD3 or BRD4, across the same cell lines in panel f.

(H) Scatter plots showing the relationship between BRD2 CRISPR score and BRD3 (left) or BRD4 (right) expression level (Log2(TPM+1)) across the same cell lines in panels f ang g. The DLD-1 cell line is highlighted in red, and Pearson correlations are indicated above.

## Notes

### Competing Interest Statement

The authors have declared no competing interest.

