## Supplementary figures and images for "BRD2 bridges TFIID and histone acetylation to promote transcriptional initiation"

### supplemental figures

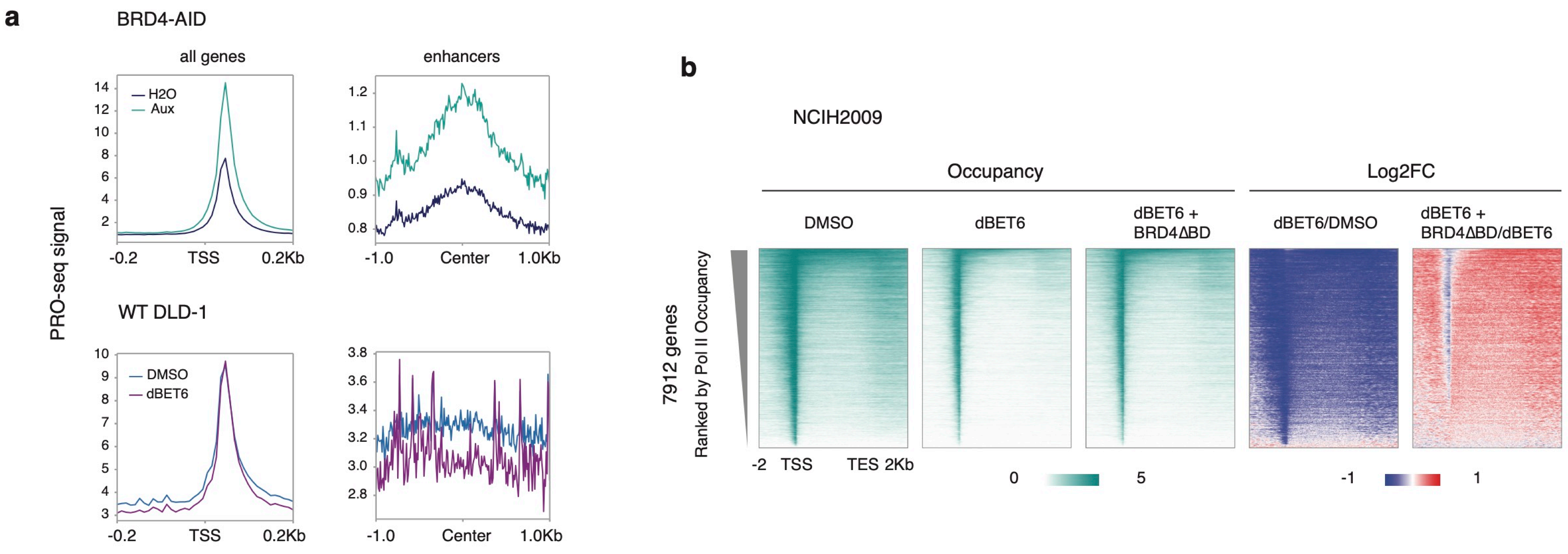

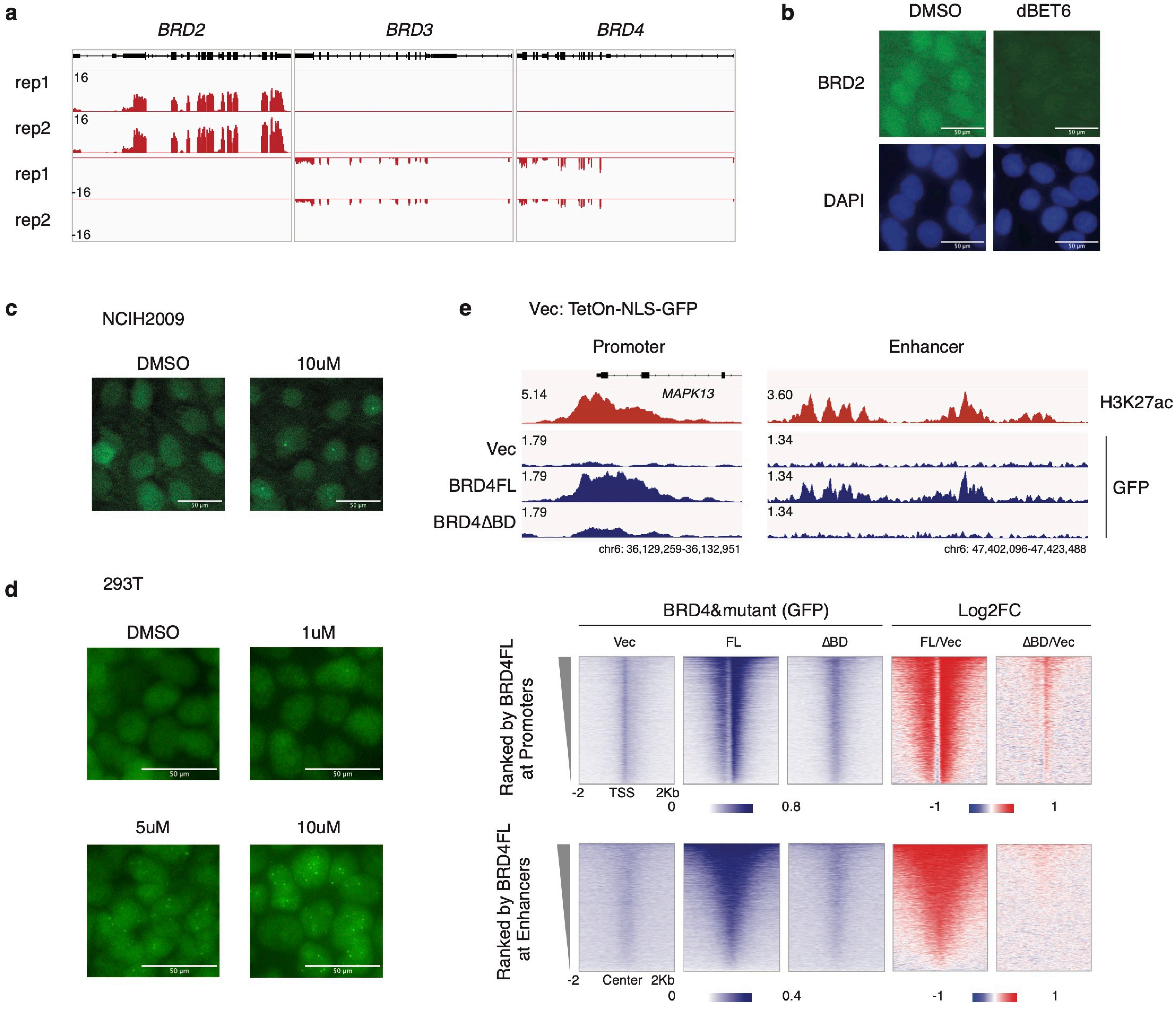

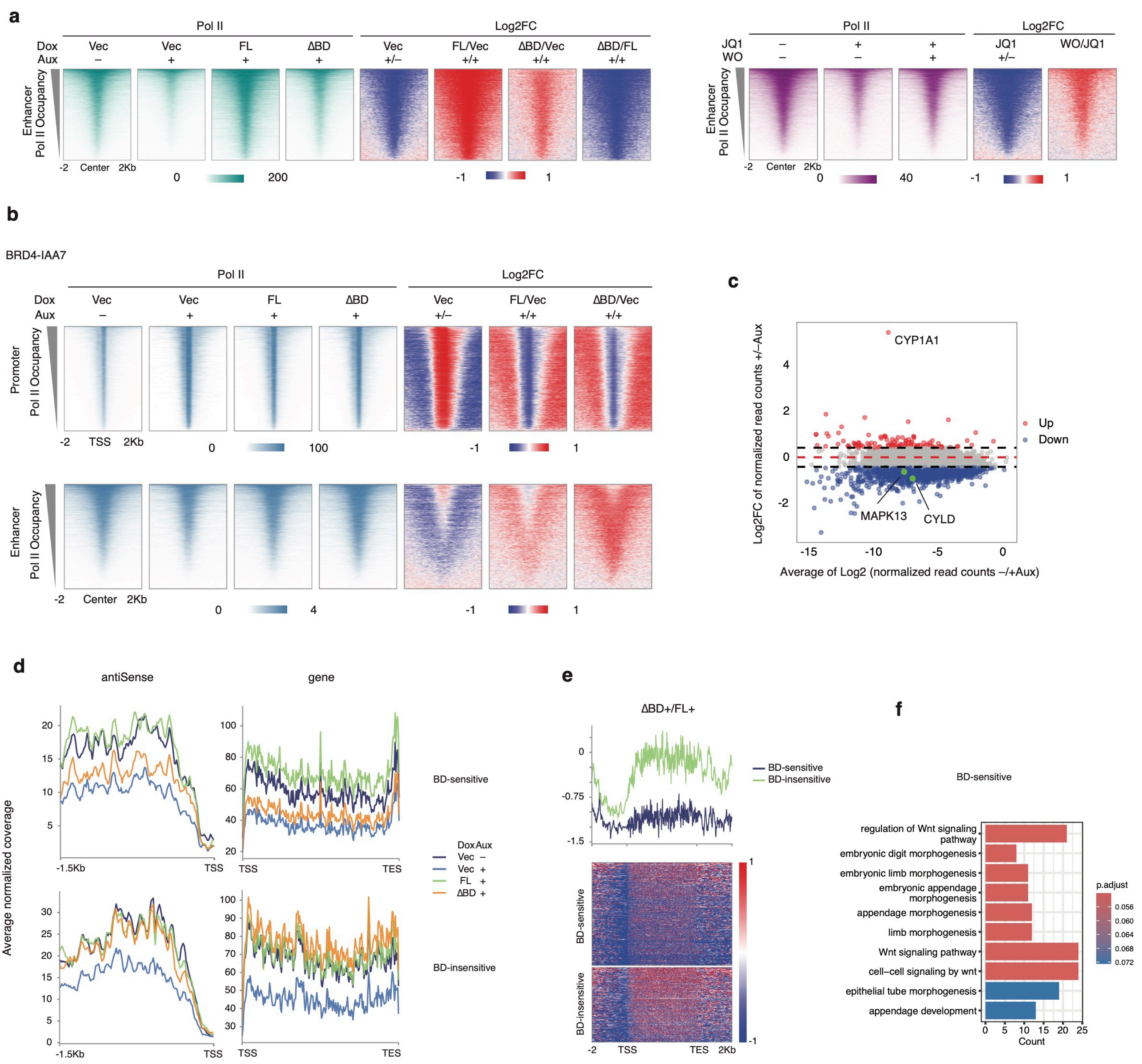

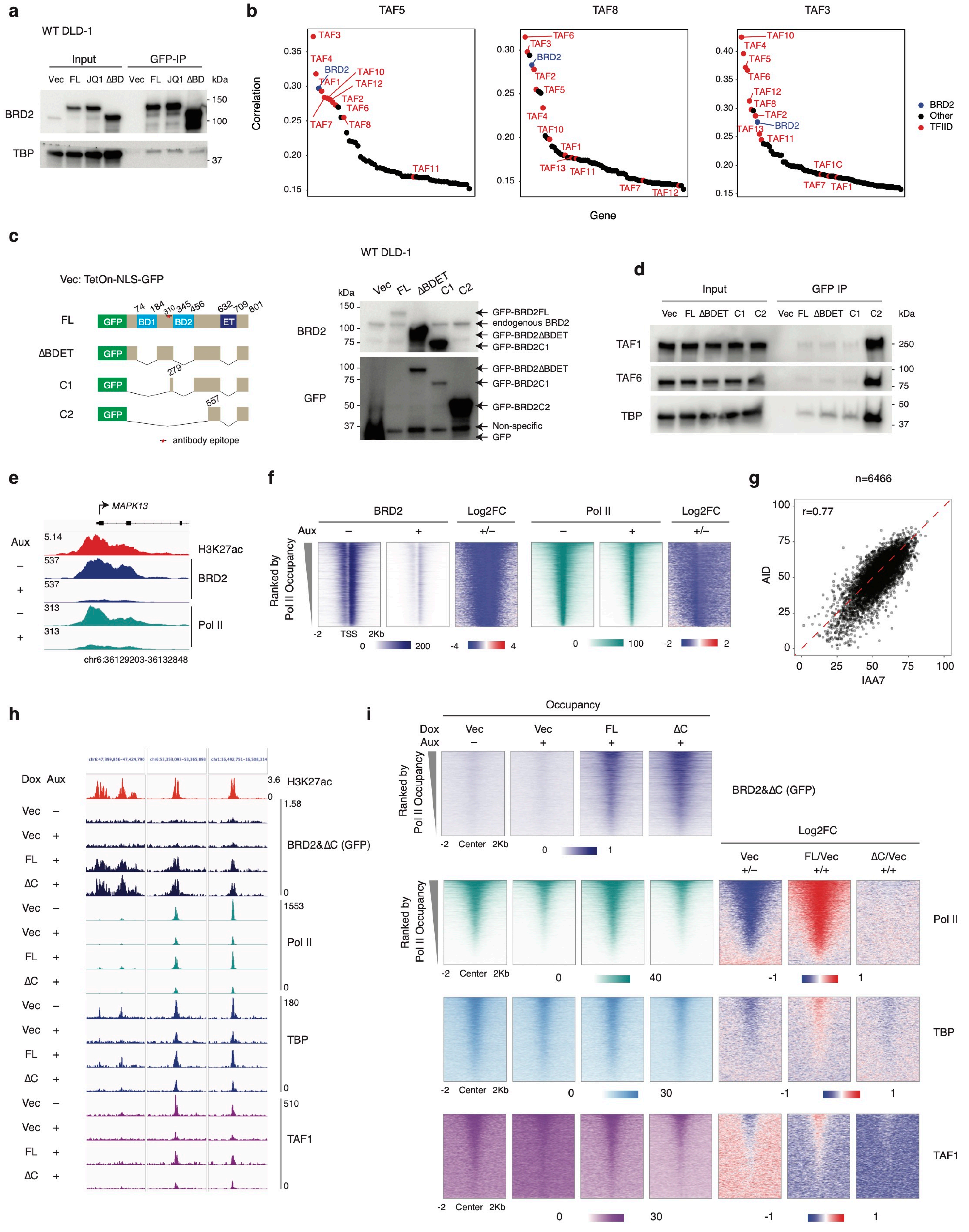

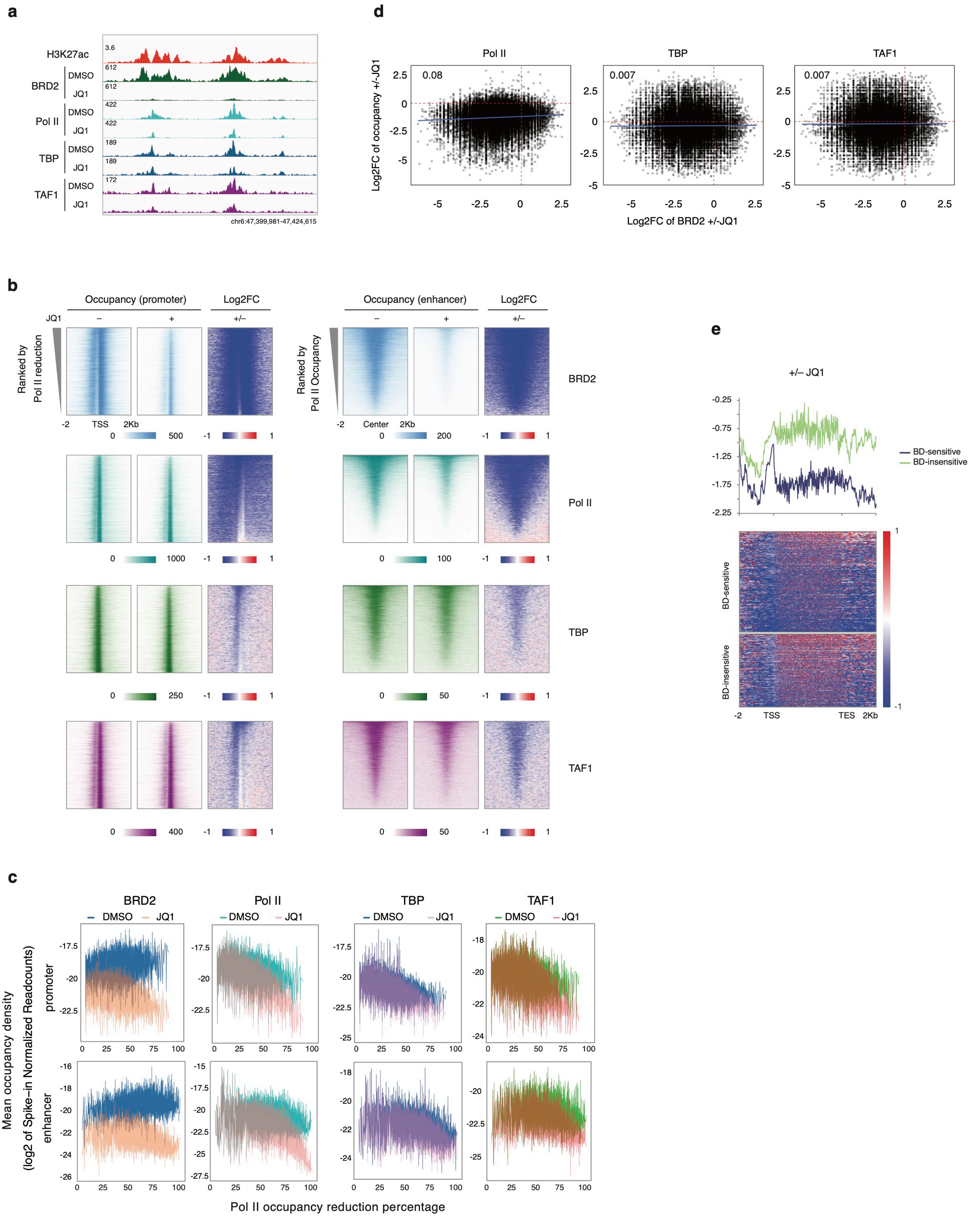

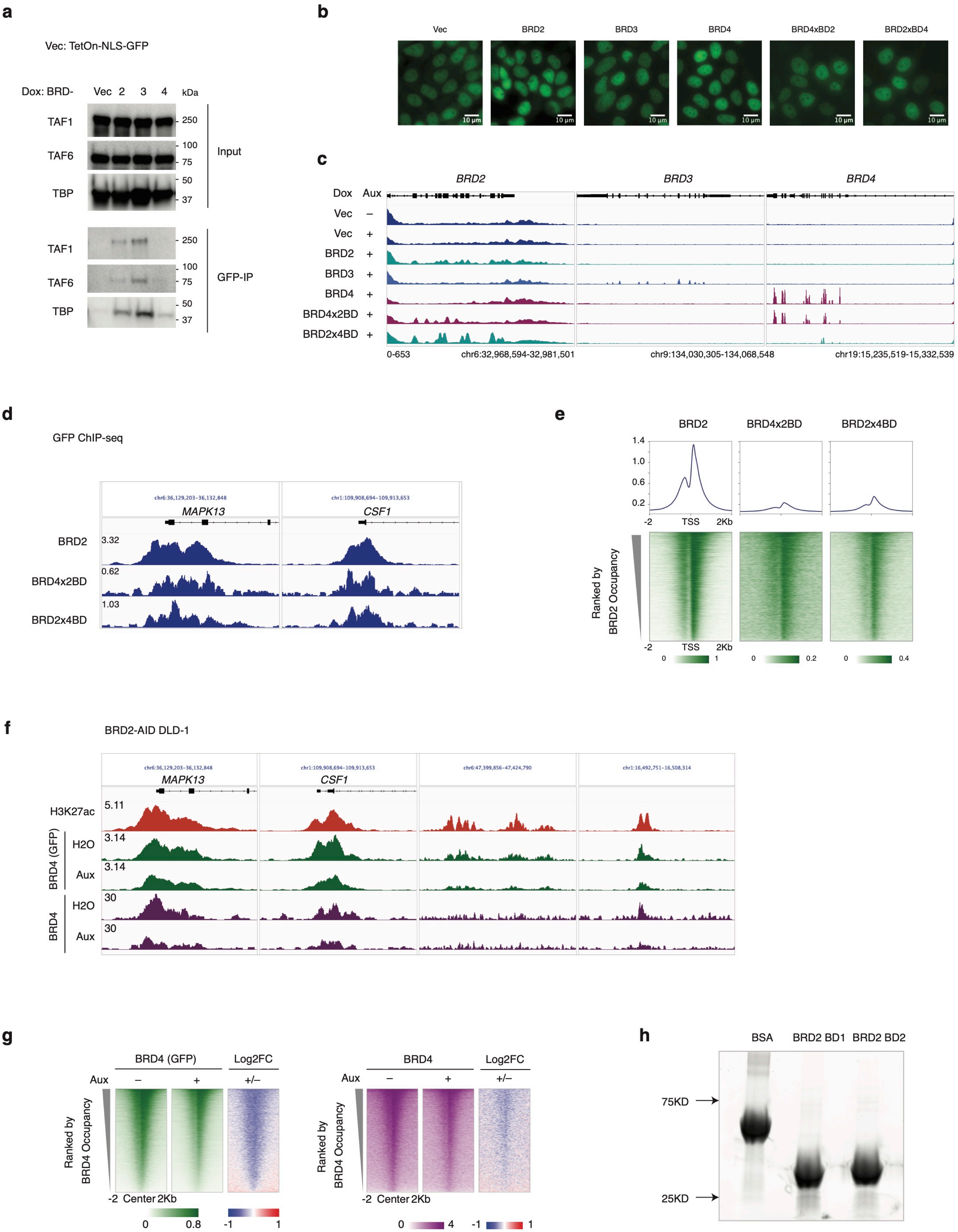

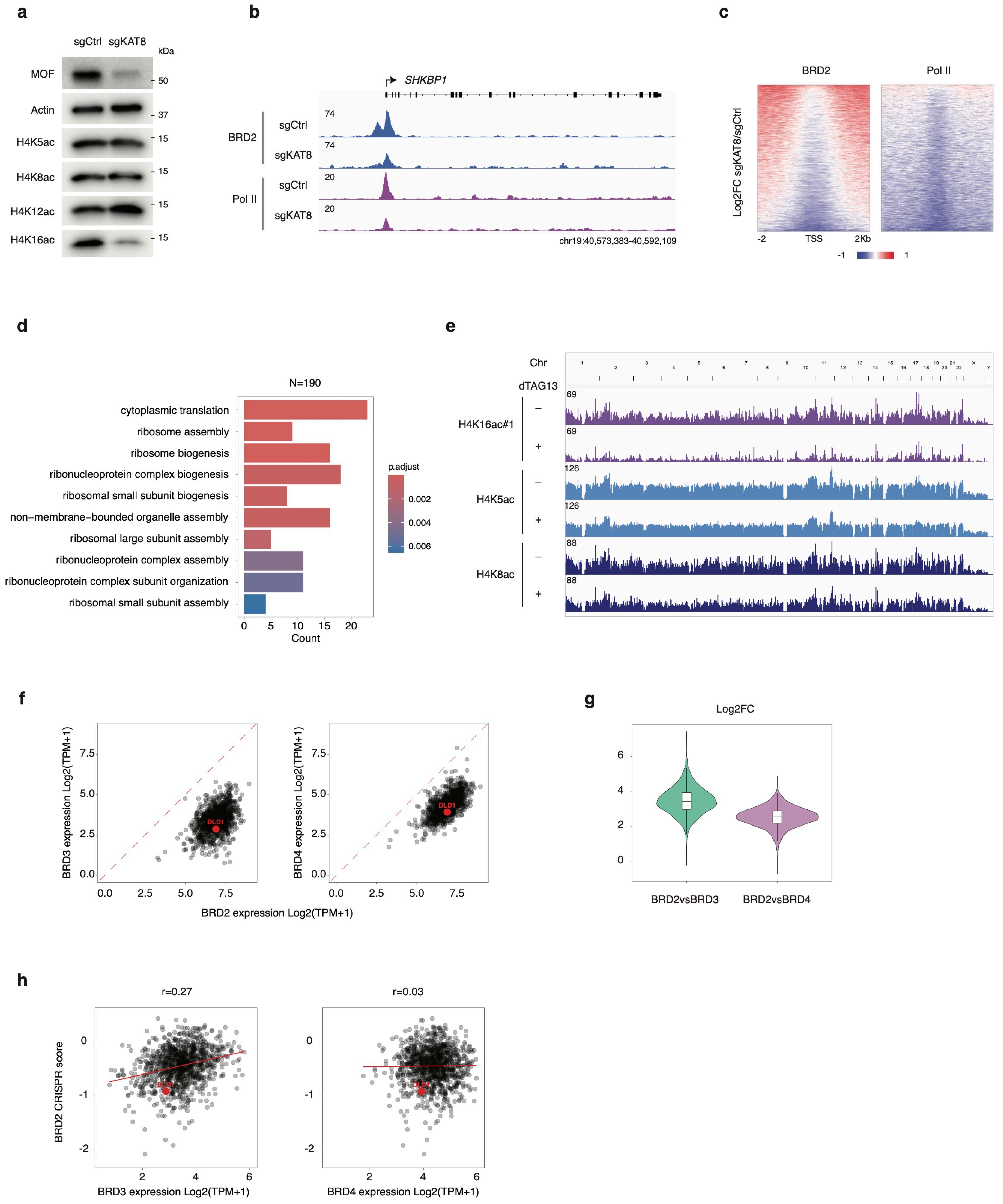
